# Low temperature triggers genome-wide hypermethylation of transposable elements and centromeres in maize

**DOI:** 10.1101/573915

**Authors:** Zeineb Achour, Johann Joets, Martine Leguilloux, Hélène Sellier, Jean-Philippe Pichon, Magalie Leveugle, Hervé Duborjal, José Caius, Véronique Brunaud, Christine Paysant-Le Roux, Tristan Mary-Huard, Catherine Giauffret, Clémentine Vitte

## Abstract

Characterizing the molecular processes developed by plants to respond to environmental cues is a major task to better understand local adaptation. DNA methylation is a chromatin mark involved in the transcriptional silencing of transposable elements (TEs) and gene expression regulation. While the molecular bases of DNA methylation regulation are now well described, involvement of DNA methylation in plant response to environmental cues remains poorly characterized. Here, using the TE-rich maize genome and analyzing methylome response to prolonged cold at the chromosome and feature scales, we investigate how genomic architecture affects methylome response to stress in a cold-sensitive genotype. Interestingly, we show that cold stress induces a genome-wide methylation increase through the hypermethylation of TE sequences and centromeres. Our work highlights a cytosine context-specific response of TE methylation that depends on TE types, chromosomal location and proximity to genes. The patterns observed can be explained by the parallel transcriptional activation of multiple DNA methylation pathways that methylate TEs in the various chromatin locations where they reside. Our results open new insights into the possible role of genome-wide DNA methylation in phenotypic response to stress.

## INTRODUCTION

Plants, as sessile organisms, must cope with constantly changing environments. When unfavorable, environmental constraints can limit normal growth and development, thus impacting plant fitness. Understanding how plants respond to environmental constraint is therefore a major task to both better understand plant local adaptation and improve agriculture. While the molecular bases of DNA methylation regulation are now well characterized, mainly in *Arabidopsis thaliana* but also increasingly in other species with more complex genomes such as rice (Tan et al. 2016), tomato (Corem et al. 2018), or maize (Li et al. 2014; Fu et al. 2018), the degree to which DNA methylation is challenged by environmental constraints remains poorly elucidated.

DNA methylation is a chromatin mark which plays an important role in various biological processes, in particular the transcriptional silencing of transposable elements (TEs). In plants, DNA methylation occurs at cytosines from CG, CHG and CHH contexts where H is A, T or G (Law and Jacobsen 2010). Several pathways regulate these different contexts in the model plant *A. thaliana*, and show preference for particular types of chromatin (Stroud et al. 2013). In TE-dense heterochromatic regions, DNA methylation is typically mediated by the action of DNA methyltransferase MET1, and chromomethylases CMT3 and CMT2 (for CG, CHG and CHH, respectively). This pathway does not necessitate active transcription but requires the chromatin remodeling factor DDM1, which facilitates access of these enzymes to heterochromatic compacted regions (Lippman et al. 2004; Zemach et al. 2013). In contrast, in gene-rich regions, DNA methylation is maintained by the action of the RNA-directed DNA methylation (RdDM) pathway (Stroud et al. 2013; Zemach et al. 2013) and requires active transcription by RNA polymerases IV and V as well as the action of DRM2, which methylates cytosines in all three contexts (Cao 2002; Onodera et al. 2005; Matzke and Mosher 2014; Cuerda-Gil and Slotkin 2016). This complex system allows for tight silencing of TEs anywhere in the genome, even in gene-rich regions where chromatin has to be kept in a relaxed form for adequate gene transcription (Zhong et al. 2012; Gent et al. 2014; Sigman and Slotkin 2016). In addition to this maintenance system, *de novo* DNA methylation can be initiated by the non-canonical RdDM pathway, which involves RNA polymerase II itself, and allows transcriptional silencing of newly inserted or deregulated TEs (Cuerda-Gil and Slotkin 2016).

Methylation sensitive amplified Polymorphism (MSAP)-based and targeted analysis of particular genomic regions have shown that DNA methylation is modified by abiotic stresses such as water deficit, metal concentration, or salinity in several species (e.g. Choi and Sano 2007; Labra et al. 2008; Verhoeven et al. 2010; Song et al. 2012). However, response of DNA methylation to abiotic constraints is still under-characterized at the whole genome scale. To date, whole-genome methylome studies have focused on the detection of small regions with differential methylation (the so-called “Differentially Methylated Regions”, or DMRs) between stressed and unstressed lots. For instance, 175 DMRs were detected following phosphate deficit in rice roots (Secco et al. 2015), 389 following cold, heat and UV treatments in maize flag leaf (Eichten and Springer 2015), or 40 following drought in *A. thaliana* leaves (Ganguly et al. 2017). So it is now clear that local DNA methylation is affected by abiotic stresses. On the other hand, observation of stress-induced global m^5^C changes through HPLC or mass spectrometry, for instance in maize (Steward et al. 2002), *A. thaliana* (Yong-Villalobos et al. 2015), *Medicago truncatula* (Yaish et al. 2018), or poplar (Lafon-Placette et al. 2018; Le Gac et al. 2018) suggests that DMRs may be only a small part of the methylome response to abiotic stress. Deciphering whether stress-induced methylation changes are regulated at the chromosome scale in link to chromatin compaction, or target specific sequences, therefore requires novel approaches. In addition, the abiotic constraints applied so far were in majority artificial, so how methylome responds to real physiological stresses encountered by plants in nature or in the field remains to be fully characterized.

Here, we investigate the methylome response of maize, a crop species with a TE-rich genome, after 3 weeks exposure to low temperatures early in development - a constraint known to induce physiological stress in this species - and propose a new strategy to analyze its impact at the whole genome scale. We show that such an exposure induces genome-wide hypermethylation in maize, which is mainly driven by hypermethylation of TEs and centromeres. These modifications affect all cytosine contexts, but CG, CHG and CHH contexts show a distinct response to cold, which intensity follows chromosome organization. The patterns observed suggest that cold enhances both RdDM and non-RdDM methylation pathways to hypermethylate TEs in all the chromatin regions where they reside. In agreement with this, a large number of genes involved in the RdDM and non-RdDM pathways are transcriptionally activated following cold. We also report 699 DMRs, which affect different types of contexts/machineries and are often associated to TE sequences. Some of these DMRs correlate with a transcription change. Altogether, these results indicate that activation of DNA methylation pathways participates in maize response to cold and suggest that tightening the transcriptional silencing of TEs is an important component of abiotic stress response to low temperature in maize, a species with high TE content.

## RESULTS

### Low temperature induces a genome-wide DNA methylation increase that mainly affects TEs

To investigate maize methylome response to abiotic stress, we exposed B73 maize plants at emergence stage to a long-term (22 days) but moderate (16 h day at 14°C; 8 h night at 10°C) low temperature treatment and compared them to control plants grown for 5 days in standard conditions, thus allowing the two sets of plants to reach similar developmental stages (Figure 1A.pdf). Plants from the stressed set showed both chlorosis and a delay in development, confirming the induction of a physiological stress (Figure 1A and B). We analyzed methylome of these plants using bisulfite-seq with 21 to 46X coverage per sample. Reads were largely (68%) uniquely aligned on the B73 genome and in vast majority (98% of these) aligned as pairs. As expected for maize, at the genome-wide scale, average cytosine methylation levels in unstressed plants were high (around 81%) for CG, medium-high (around 68%) for CHG and low (around 1%) for CHH contexts (Supplemental_Table_S1.pdf). Interestingly, average methylation levels were a few percent higher in stressed than unstressed plants, overall and for each context separately, and particularly for CHGs (71% *vs.* 68%). Stressed samples also showed higher number of methylated cytosines for all three contexts, and was significant for CHG (Student’s test, p-value 0.03, Supplemental_Table_S1.pdf). Altogether, our results indicate that low temperature induces a global genome hypermethylation in the B73 cold-sensitive inbred line.

**Figure 1:**
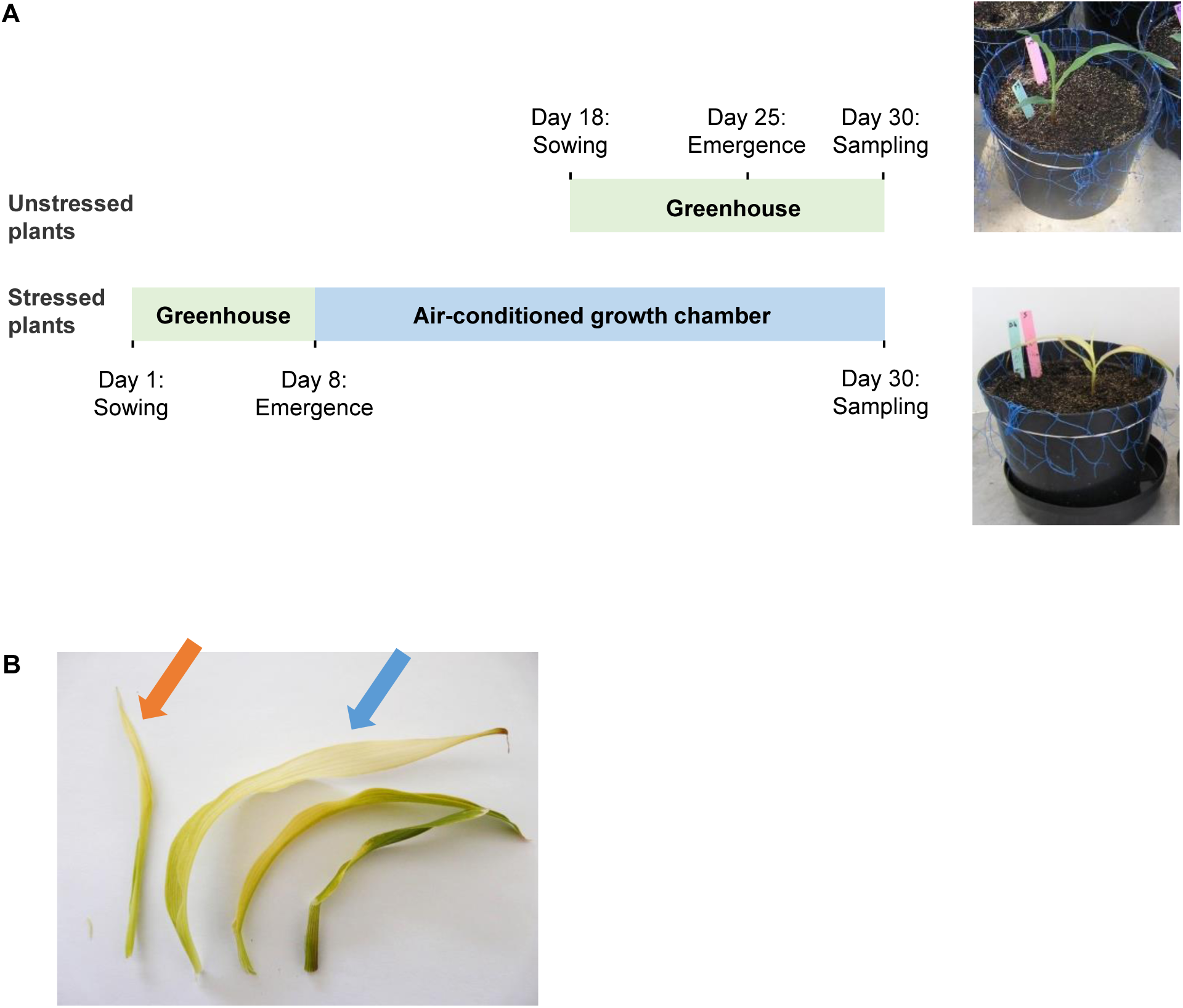
Experimental setting, plant phenotype and leaf sampling. **A.** Experimental setting with phenotype of the plants. At emergence stage, plants were placed either in low temperature conditions (air conditioned chamber, 16h day at 14°C; 8h night at 10°C) or in standard conditions (greenhouse, 16h day at 24°C; 8h night at 18°C) leading to two sets of plants referred to as ‘‘stressed’’ and ‘‘unstressed’’. Day length of the air-conditioned growth chamber was synchronized to this of the greenhouse. Due to an impact of low temperature on plant development, the unstressed set was sown 18 days after the stressed plants were sown, thus allowing stressed and unstressed plants to reach a similar developmental stage at the end of the experiment. After exposure to low temperature, stressed plants show important chlorosis. **B**. Detail of the leaves sampled. The last visible leaf (orange arrow) and the one before visible leaf (blue arrow) of 3 individual plants of each set was sampled for methylome and transcriptome analysis, respectively.

To investigate whether this hypermethylation was homogeneous across chromosomes, we first analyzed the average DNA methylation difference (average methylation value of unstressed plants minus average methylation value of control plants) across chromosomes using 100 kb bins, and showed that hypermethylation is observed throughout chromosomes (results for chromosome 10 are illustrated in Figure 2A; for the other chromosomes, see Supplemental_Fig_S1.pdf).

**Figure 2:**
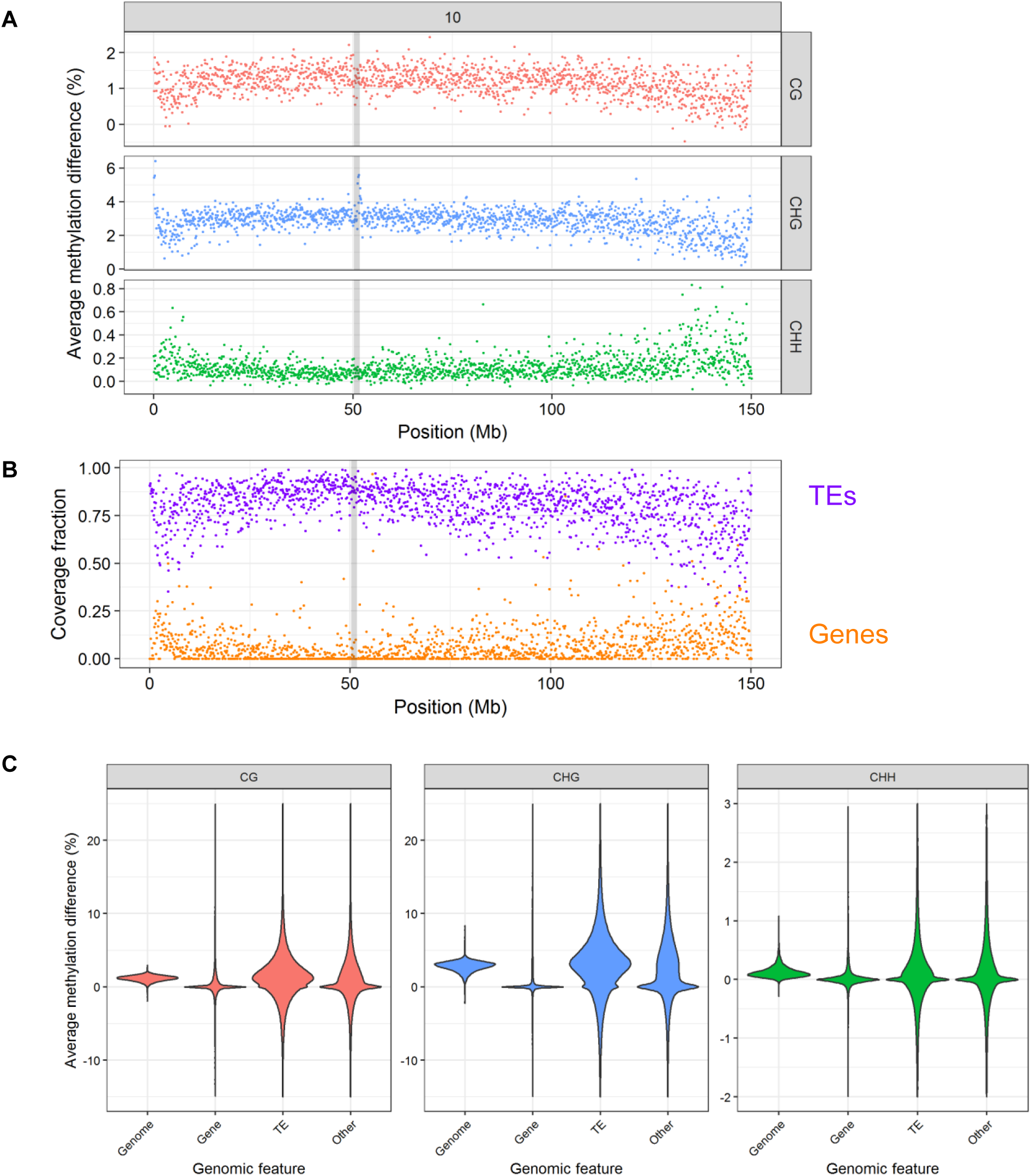
Methylation response to low temperature across chromosomes and for the different genomic features. **A.** Chromosome 10 methylation landscape showing average difference between methylation levels of stressed and unstressed plants over 100 kb bins, for each of the three cytosine contexts. Contexts are shown on the right, in grey panels. Positions are in Mb from tip of chromosome 10. Centromere position is highlighted by a grey box. **B.** Chromosome 10 annotation landscape showing percent of coverage of each 100 kb bin by TE (purple) and gene (orange) bases. Positions are in Mb from tip of chromosome 10. Centromere position is highlighted by a grey box. Profiles of the other chromosomes are presented in Supplemental_Fig_S1.pdf. **C.** Violin plot of methylation response for the different genomic features. Cytosine contexts are shown on top.

We found that methylation landcapes are specific of each cytosine context, and vary in relation to the density of genes and TEs: for both CG and CHG contexts, methylation difference is highest in TE-rich regions, while difference at CHH is highest in gene-rich regions (Figure 2A and B, Supplemental_Fig_S1.pdf). To test whether this effect was due to targeting of specific features or to chromosomal location, we then analyzed the DNA methylation patterns of genes, TEs and non-genic non-TE sequences (referred to as “Other”). While genes were almost invariable, TEs showed a large enrichment for methylation change in all three cytosine contexts, and Other sequences showed an intermediate pattern (Figure 2C). Although some hypomethylation cases were observed, hypermethylation cases were by far more numerous and with a higher methylation change, thus leading to observation of a global hypermethylation at the chromosome scale.

We then tested whether this feature-based effect was combined with a chromosomal localization effect. For CG and CHG, levels of variation at genes, TE and Other features did not vary across chromosome (Supplemental_Fig_S2.pdf), highlighting that chromosomal bell-shaped landscape pattern is due to the higher density of TEs in the middle of the chromosome, rather than targeting of specific TEs in particular chromosomal regions. For CHH however, while levels of variation at genes and Other features were largely similar across chromosomes, for TEs, cases with the highest levels of variation were found predominantly in gene-rich regions, highlighting an effect of chromosome localization for this context (Figure 3A, Supplemental_Fig_S2.pdf). To further investigate this effect, we analyzed the methylation response of TEs located within 5kb of genes. As shown in Figure 3B, while CG and CHG methylation was increased for all types of TEs, a different behavior was observed for CHH between TEs located within genes or in their close proximity. TEs overlapping genes showed almost no mCHH variation. In contrast, TEs immediately flanking genes showed the highest variation, and the variation level gradually decreased with distance to the closest gene, reaching basal levels around 2 kb from the gene border (Figure 3B). This highlights that not only proximity to gene, but the distance itself, is important in the TE CHH methylation response to low temperature. A difference between genic TEs and TEs located nearby genes was also observed for CHG but was more limited, as genic TEs showed a certain extent of methylation variation (Figure 3B, Supplemental_Fig_S3.pdf). In the CHG case, however, distance to the nearest gene had only a subtle effect on the TE methylation variation observed, with a slight increase up to 1kb from the gene. Interestingly, both the inside/nearby difference and the distance effect observed for CHH methylation was particularly marked for three out of the four DNA transposon super-families, as well as for unclassified LTR retrotransposons (RLX), while almost absent for CACTA transposons (DTC) and for LTR retrotransposons of the *Gypsy* (RLG) and *Copia* (RLC) super-families (Figure 3C). LINE response was difficult to assess due to limited number of cases. For all five TE superfamilies showing a methylation response, the effect was more marked for copies located in the upstream regions of genes (Figure 3C, Supplemental_Fig_S4.pdf and Supplemental_Fig_S5.pdf). Altogether, these results highlight the impact of low temperature on the remodeling of TE methylation, with an effect of chromosome location, distance to closest gene, and TE type.

**Figure 3:**
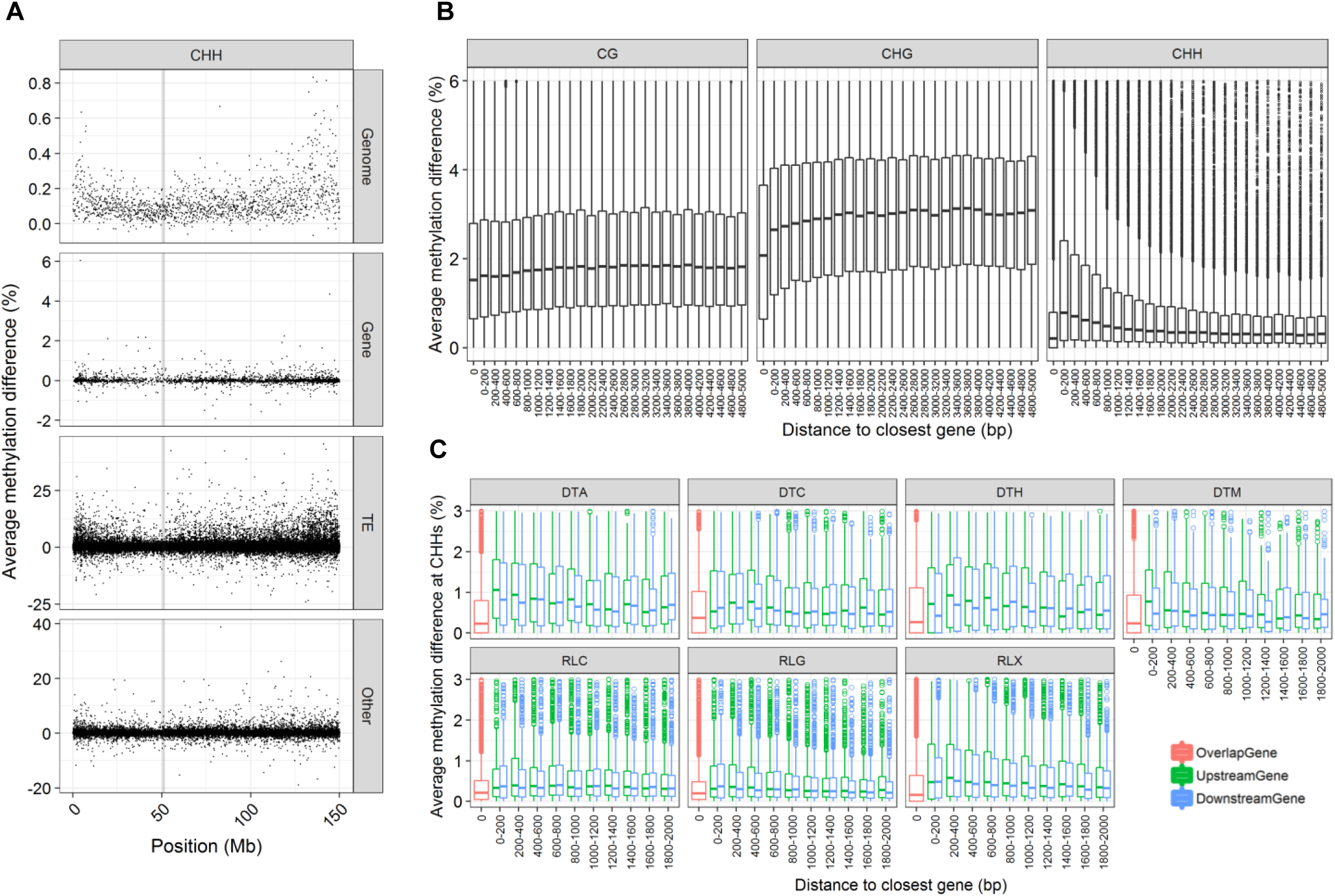
Impact of gene distance on TE methylation response. **A.** CHH methylation difference landscape for each category of genomic feature, example of chromosome 10. Genomic feature type is shown in right panels. Each dot corresponds to one genomic region. For genome, regions are all of 100 kb. For genomic features, sizes are variable. Centromere position is highlighted by a grey box. Landscapes for all chromosomes and all contexts are presented in Supplemental_Fig_S2.pdf **B.** Distribution of methylation difference in TEs for CG, CHG and CHH contexts, with respect to distance to closest gene. Only hypermethylated cases are shown, see Supplemental_Fig_S3.pdf for hypomethylated cases. **C.** CHH methylation difference for TEs located upstream (green), downstream (blue), or overlapping (red) genes. Each panel shows one TE class. DTT, RIL and RIX are not represented due to too low sample size. Corresponding graphics for CG, CHG, CHH of hypermethylated and hypomethylated cases are presented in Supplemental_Fig_S4.pdf and Supplemental_Fig_S5.pdf, respectively.

When analyzing the methylation chromosomal landscapes, we also noticed regions with very high CHG increase (>5% over several bins of 100 kb) at centromeres, as well as at particular positions of chromosomes 7 and 10 (Figure 2A, Figure 4A and B, Table 1). In these regions, all types of features showed a higher methylation change than the rest of the genome (Figure 4C), suggesting a targeting of these particular genomic regions rather than particular sequences. Because centromeres are known to harbor the particular histone variant CENH3, we then compared the position of our CHG peaks with these of CENH3 ChIP-seq experiments (Zhao et al. 2015). We found a clear association between the two, highlighting that cold induces a CHG hypermerthylation at CENH3-containing regions.

**Table 1:** Segmentation results of CHG average methylation difference between stressed and unstressed plants

| Chromosome | Region type | Segmentation start | Segmentation end | Number of datapoints in segment | Mean methylation difference in segment |
| --- | --- | --- | --- | --- | --- |
| 2 | Centromere | 93.4 | 94.6 | 13 | 4.64 |
| 3 | Centromere | 99.8 | 100.8 | 11 | 4.73 |
| 5 | Centromere | 102.1 | 103.2 | 12 | 4.67 |
| 6 | Centromere | 49.6 | 50.1 | 6 | 4.80 |
| 8 | Centromere | 49.8 | 51.2 | 13 | 5.10 |
| 9 | Centromere | 51.5 | 52.3 | 9 | 4.75 |
| 10 | Centromere | 51.2 | 52.1 | 10 | 4.48 |
| 7 | Additional | 22.7 | 22.9 | 3 | 6.63 |
| 10 | Additional | 0.1 | 0.4 | 4 | 5.25 |

**Figure 4:**
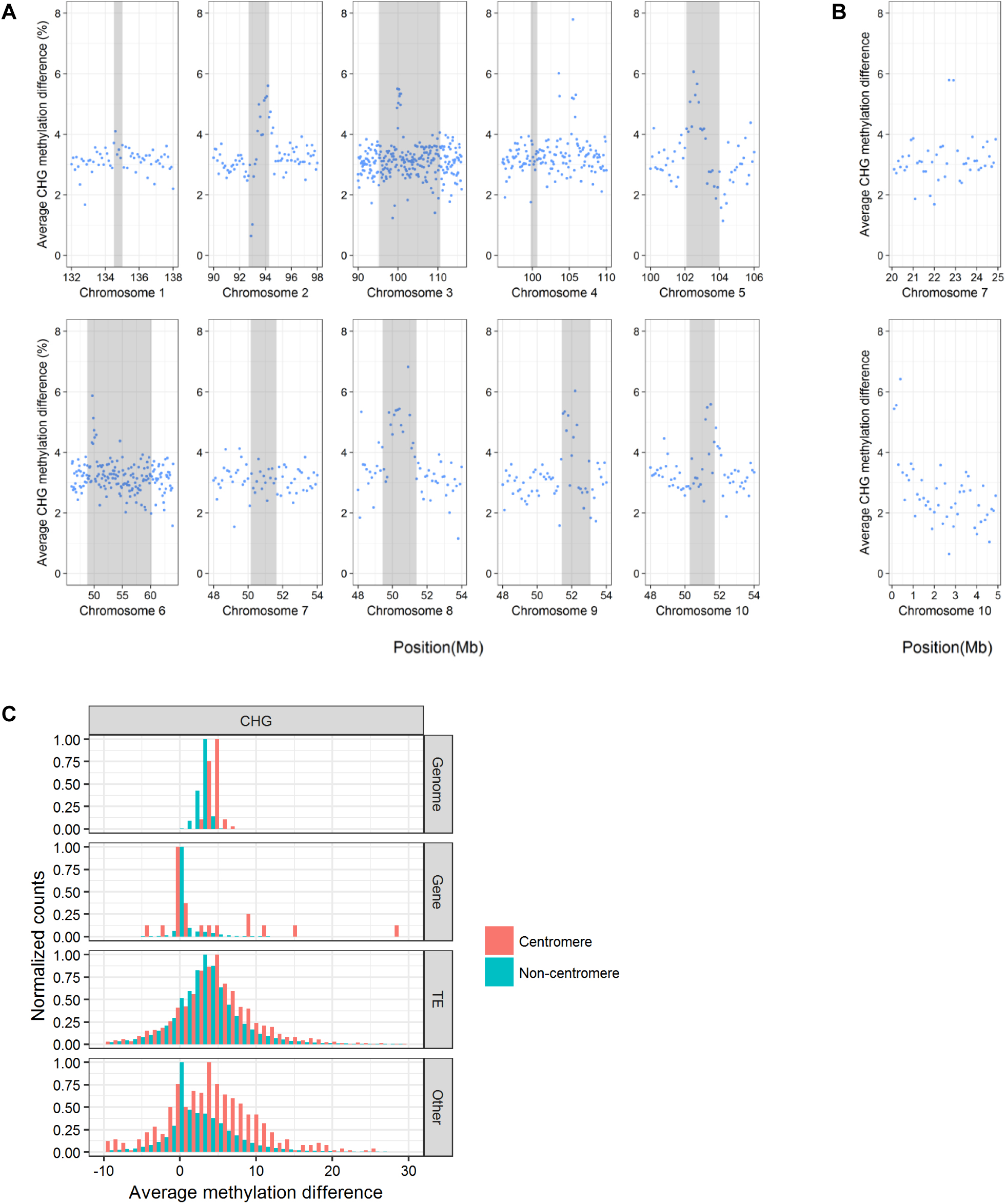
CHG methylation change at centromeres. **A.** CHG methylation peaks at centromeres. Each dot corresponds to a 100 kb bin. Position of centromeres are shown in grey. Positions are in Mb from chromosome tip. **B.** CHG methylation peaks observed at other regions on chromosomes 7 and 10. Each dot corresponds to a 100 kb window. Positions are in Mb from chromosome tip. **C**. Average CHG methylation difference for centromeric (red) and non-centromeric (green) regions. Feature types are shown on the right. Counts are normalized to maximum value of each category.

### Local Differentially Methylated Regions (DMRs) are associated with TEs

The chromosome-and feature-centered approaches allowed us to decipher the large-scale DNA methylation changes induced by low temperature in maize. To investigate the impact of this treatment at a more local scale, we also detected Differentially Methylated Regions (DMRs) between stressed and unstressed plants, using a combination of 200, 500 and 1 kb windows and requiring a minimum methylation change of 10% between stressed and unstressed lots, and a minimum of 5 Differentially Methylated Cytosines (DMCs) with at least 10% change within the DMR (see Material and Methods for details). Considering the low levels of CHH methylation, we focused only on CG and CHG DMRs. With these criteria, we detected a total of 750 DMRs, among which 354 and 396 were hypermethylated or hypomethylated in stressed plants, respectively (Supplemental_Table_S2.pdf). Characteristics of these DMRs are depicted in Supplemental_Table_S2.pdf, and Supplemental_Fig_S6.pdf to Supplemental_Fig_S9.pdf. Hypermethylated DMRs were slightly longer and showed higher DNA methylation variation than hypomethylated ones.

To get insights into possible underlying mechanisms, we then classified these DMRs based on their variation in either CG (“CG only”), CHG (“CHG only”) or variation in both CG and CHG (“CG + CHG”) using a 2-component Gaussian mixture model (Supplemental_Fig_S10.pdf and Supplemental_Fig_S11.pdf). After removing the redundancy among CG+CHG category, we finally characterized a total of 377 hypomethylated DMRs and 322 hypermethylated DMRs. Most (52%) classified DMRs vary both in CG and CHG, 32 % vary only in CHG, and a smallest fraction (7%) varies only in CG (Supplemental_Table_S3.pdf). CGonly DMRs show the smallest variation both for hyper-and hypo-methylated cases (on average, 18.5 and 16.3%, respectively, Supplemental_Fig_S10.pdf and Supplemental_Fig_S11.pdf), exhibit a wide range of initial methylation levels (from 0 to 68%), and are devoid of methylation at CHGs and CHHs (Figure 5A, left). They are enriched inside genes, and are associated to genic TEs but are also abundant away from genic TEs (Figure 5A, right). Therefore, they are likely generated by the genic CG methylation maintenance. CHGonly DMRs show a similar range of initial methylation levels, and absence of CHH methylation, but their CG methylation levels are close to 100% (Figure 5B, left). They are enriched inside genes and highly associated with genic TEs (enriched within or near genic TEs, and depleted away from genic TEs). Hence, CHG DMRs are likely generated by genic TE CHG methylation maintenance. CG+CHG DMRs show variation in all three contexts (Figure 5C), and are enriched close to TEs, independently of their proximity to genes. They therefore likely originate from RdDM methylation maintenance at TE borders. Therefore, the large majority of the DMRs found is associated to TE methylation regulation. We found similar patterns of annotation for both hyper-and hypomethylated DMR cases (Figure 5, Supplemental_Fig_S12.pdf), suggesting that low temperature can induce local enlargement or contraction of methylation at TE boundaries.

**Figure 5:**
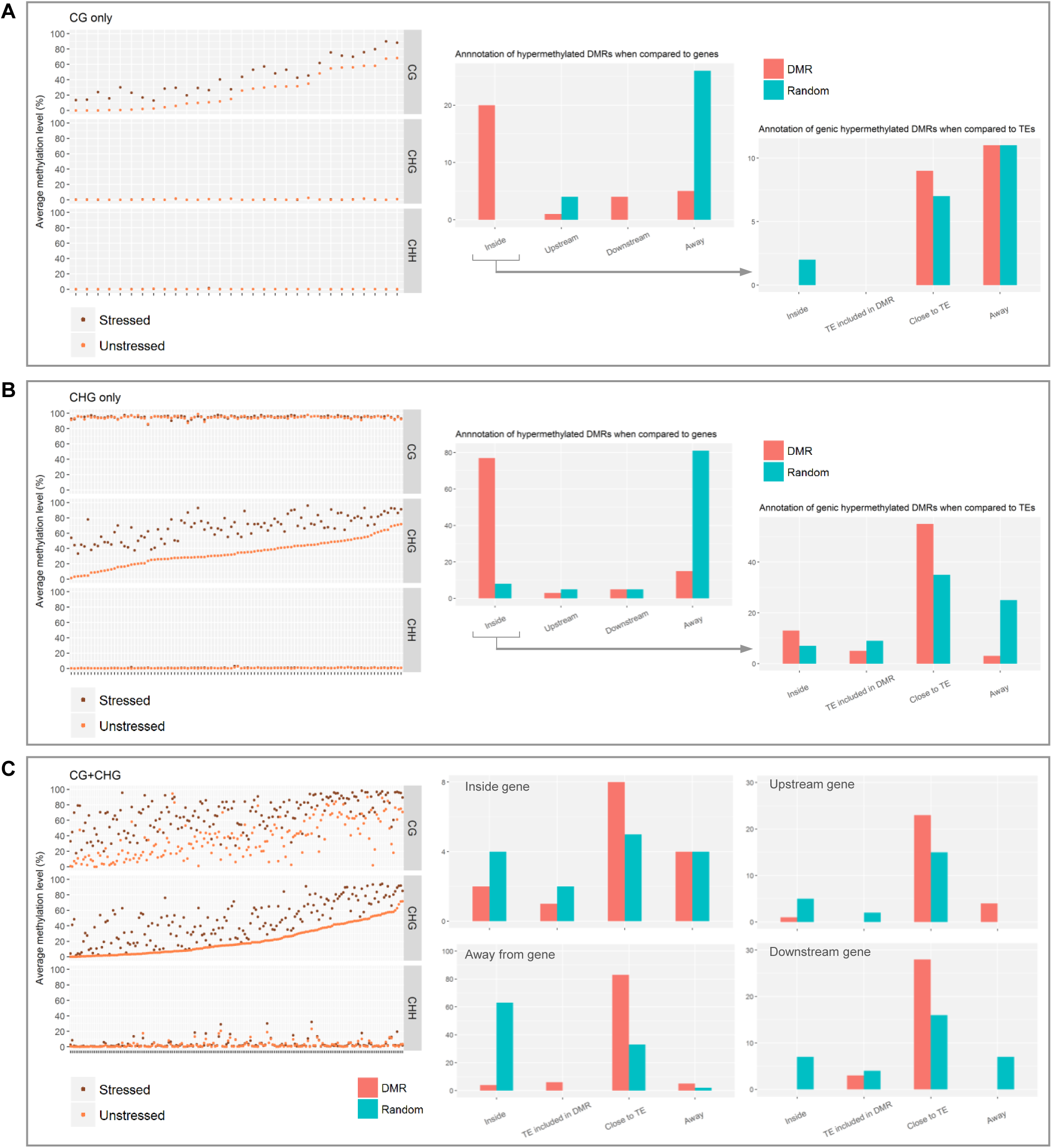
Characteristics of hypermethylated DMRs. **A.** Caracteristics of CG only DMRs. Left: methylation levels in stressed and unstressed plants. Each position of the x axis corresponds to one DMR, to which two dots correspond, with average values in stressed (brown) and unstressed (orange) plants. Middle: position relative to genes as compared to random regions. For category with largest enrichment (DMRs located inside genes), position relative to TEs is given on the right panel, as compared a random set of regions located inside genes. **B.** Caracteristics of CHG only DMRs. Same legend as A. **C**. Caracteristics of CG+CHG DMRs. Left: same legend as panels A and B. Right four panels: DMR position relative to TEs as compared to random regions, after subdivision of DMRs as compared to distance to genes. Distance thresholds are 2 kb and 500 bp, for genes and TEs, respectively. Characteristics of hypomethylated DMRs are presented in Supplemental_Fig_S12.pdf.

### Cold-induced transcriptional changes highlight a physiological stress and perturbation of the RdDM and non RdDM methylation maintenance pathways

To get further insights into maize molecular response to prolonged low temperature, we analyzed the transcriptomic changes between stressed and unstressed plants. Strand specific 2×150 bases paired-end mRNA-seq data generated from the same plants as those used for methylome studies were mapped onto B73 reference genome sequence, with 90% of unambiguously mapped pairs. Statistical analysis with a Benjamini-Hochberg procedure (FDR<0.05) revealed a total of 9030 differentially expressed genes between stressed and unstressed plants, 4621 and 4409 being up-and down-expressed in stressed plants, respectively. Differentially expressed genes showed highly significant enrichment for particular gene ontologies classes: downregulated genes showed an enrichment in photosynthesis (40/117 genes, pvalue= 3.36e^-09^), lipid biosynthesis (75/313, pvalue= 8.98e^-09^), oxydo-reduction (319/1929 genes, pvalue= 6.82e^-07^), cellulose biosynthesis (18/50 genes, pvalue=4.59e^-06^), while upregulated genes were enriched in translation (294/1213 genes, pvalue<1.00e^-20^), RNA processing (80/297 genes, pvalue= 2.76e^-14^), ribosome biogenesis (30/68 genes, pvalue= 2.76e^-14^), RNA modification (24/53 genes, pvalue= 2.47e^-10^), and tRNA metabolic process (52/203 genes, pvalue= 2.95e^-09^). These results highlight that the low temperature treatment to which plants were exposed induced a physiological stress at the cellular level, and reflect the phenotype observed for stressed plants. Interestingly, several pathways generating various components of cell wall such as cellulose, lipids and membrane transporters are perturbated in our experiment, highlighting the large impact of low temperature on cell wall structure and function.

Finally, to further investigate origin of the methylome response observed, we specifically analyzed the expression pattern of a series of 80 genes described in the literature as involved in DNA methylation regulation (Supplemental_Table_S4.pdf). A total of 23 genes (29%) showed significant differences between stressed and unstressed plants, among which 7 were down-regulated and 16 were up-regulated in stressed plants (Table 2). Interestingly, 8 of the 16 up-expressed genes are involved in the canonical RdDM pathway. They encode RNA polymerases IV and V components (NRP(B/D/E)6a, NRP(D/E)7) and NRP(D/E)10c), interacting factors (IWR1/DMS4/RDM4, SHH2c and MOP1), as well as AGO proteins (AGO104 and AGO105/AGO119). Genes involved in CG maintenance such as *Zmet8/Zmet1/Zmet1b* and in CHG maintenance such as *Kyp* also showed an expression increase in stressed plants, as well as the chromatin remodeling factor *Chr101/DDM1a*. Altogether, these results suggest the parallel activation of the RdDM and the non-RdDM methylation maintenance pathways, thus supporting an increase of methylation maintenance. Surprisingly, we observed an expression decrease for methyltransferase genes *Zmet2/Dmt102/ZmCMT1/ZmMet2a* and *Zmet3/Dmt103/ZmDRM1/ZmMet3b*, suggesting a decoupling between activation of the pathways and action of the methyltransferases *per se*. The expression increase observed for *Zmet6/Dmt106/ZmDRML/ZmMet3c* also suggests a possible transcriptional replacement of *Zmet3/Dmt103/ZmDRM1/ZmMet3b* by *Zmet6/Dmt106/ZmDRML/ZmMet3c* in response to cold.

**Table 2:**
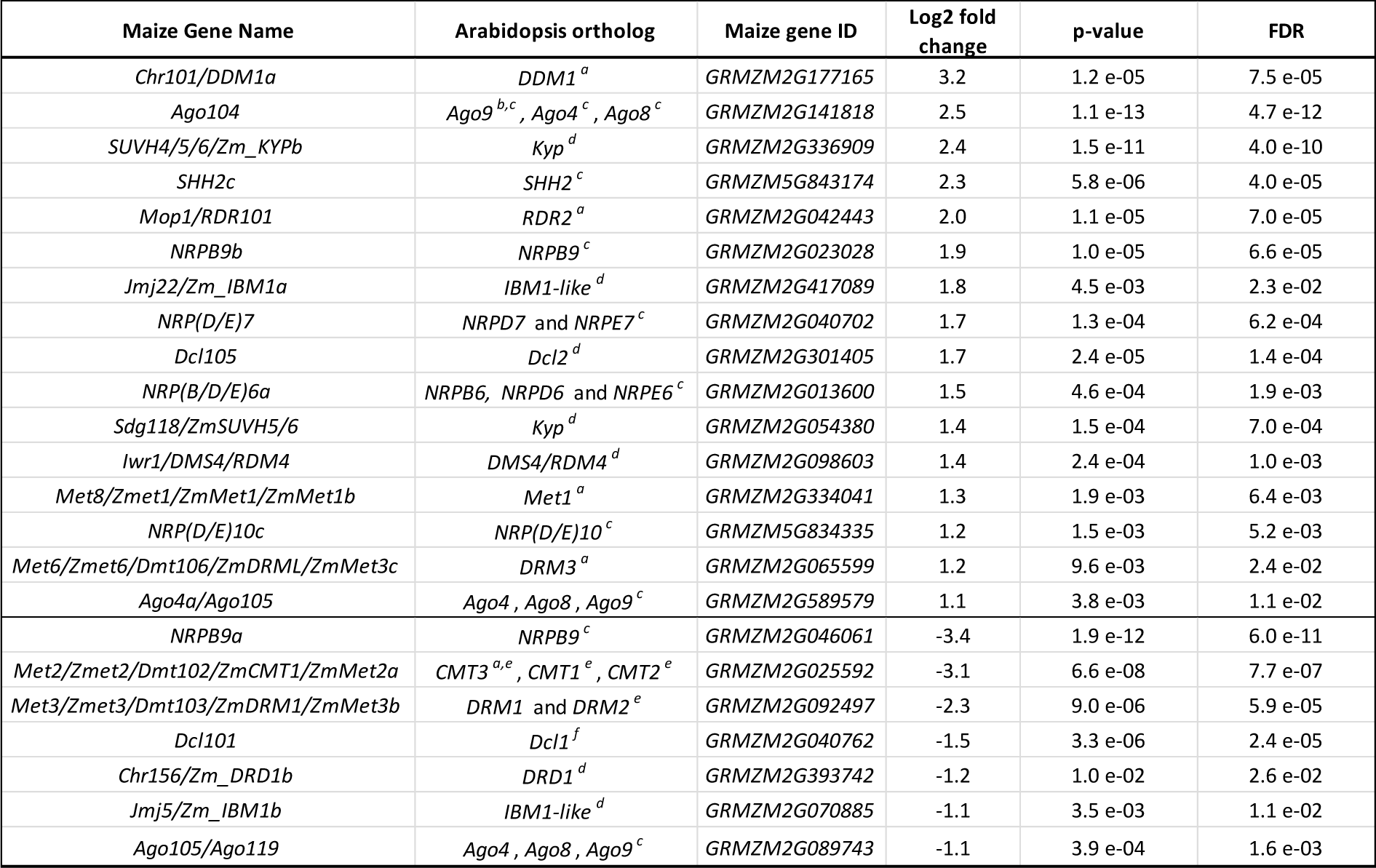
Differentially expressed genes involved in DNA methylation regulation

Behavior of the *de novo* RdDM machinery was less clear, with paralogous genes coding for polymerase II subunit NRPB9 showing an expression increase (NRPB9a) or a decrease (NRPB9a), and *Dcl105* and *Dcl101* showing a contrasted response, namely an increase for *Dcl105* and a decrease for *Dcl101* (Table 2).

Genes are ranked based on Log2FoldChange between the low temperature and standard conditions. Normalized counts: average count values for unstressed plants, after normalization by TMM and gene size. Log2FoldChange are positive when upregulated in stressed plants, and negative when downregulated in stressed plants. FDR: False discovery rate based on Benjamini-Hochberg correction. Information on Arabidopsis orthologs is extracted from: a: Li et al. 2014; b: Singh et al. 2011; c: Haag et al. 2014; d: *Arabidopsis thaliana* best hit on MaizeGDB e: Candaele et al. 2014, f: Petsch et al. 2015.

### DMRs associate with genes involved in stress response, some showing a transcriptional change

Among the 699 DMRs, a total of 418 (59%, 218 hypomethylated cases and 200 hypermethylated cases) were found located in or near a protein coding gene (located within a 2 kb distance). To get insights into possible methylation-based gene regulation, we further analyzed the corresponding genes. First, we tested whether genes with a DMR located within a 2 kb distance were enriched in particular functions. GO enrichment analysis of the 200 hypermethylated DMRs revealed enrichment in protein arginylation (1/1 gene, pvalue=4.49e^-03^), fucose metabolic process (1/2 genes, pvalue=8.96e^-03^), double-strand break repair *via* nonhomology end joining (1/2, pvalue=8.96e^-03^). This of the 218 hypomethylated DMRs revealed enrichment in actin cytoskeleton organization (3/62 genes, pvalue=4.69e^-03^), and double-strand break repair *via* homologous recombination (1/1 gene, pvalue=5.44e^-03^). Interestingly, a total of 28 genes involved in stress response (GO 0006950) had associated DMRs (16 hypermethylated, 12 hypomethylated). Among these, 6 genes (21%) also showed a change in transcription. These encode a heat-shock protein (*GRMZM2G024668*), a peroxidase (*GRMZM2G117706*), a nucleosome-binding factor (*GRMZM2G128176*), an ATP-dependent DNA helicase (*GRMZM2G137968*), a sphingolipid transporter (*GRMZM2G062024*), two proteins involved in cell division control (*GRMZM2G149994* and *GRMZM2G166718*), three proteins involved in DNA repair (*GRMZM2G016602, GRMZM2G030128* and *GRMZM2G035417*), a protein involved in cell wall composition (*GRMZM2G005562*), and a protein involved in the blockage of mitosis (*GRMZM2G098637*). Function of these genes are consistent with the phenotypic response observed in stressed plants, which show a delay in development, leaf chlorosis and cell wall perturbation, suggesting a role of methylation change in transcriptional and phenotypic response to cold. DMRs associated with a transcriptional variation in stress-related genes showed a slight enrichment for location inside genes (9/10 cases, *vs*. 21/28 cases for all DMRs associated with stress-related genes). The mechanistic link between methylation and transcription was nevertheless difficult to assess, with no enrichment in hyper-or hypomethylated cases, and no clear link between orientation of methylation and transcription (6 correlated cases *vs*. 4 anticorrelated cases).

### Transcriptional changes of genes associated with DMRs vary depending on DMR location

To get further insights in to possible link between methylation and transcription, we then extended this analysis to all classified DMRs located within 2 kb of a gene (377 DMRs, including 192 hyper-and 185 hypo-methylated cases). We found that 117 of these DMRs (31%) are close to a differentially expressed gene (Supplemental_Table_S5.pdf). Similar proportions were observed for the subset of DMRs located inside genes (75/223, 34%), as well as for DMRs located downstream genes (29/82 cases, 36%). In contrast, only 19% (13/70 cases) of genes with a DMR upstream showed a transcriptional change, suggesting that the impact of methylation variation is less important when located upstream genes. Interestingly, for DMRs located upstream or downstream genes, transcriptional changes was observed more often for hypomethylated cases than hypermethylated cases (24% vs. 12% for upstream DMRs, and 50% vs. 24% for downstream DMRs), suggesting that a decrease in methylation in regulatory regions has a stronger effect on transcription.

DMRs located inside genes are associated with an increase in transcription. But this is independent of the methylation change observed, with 63% of hypomethylated DMRs and 68% of hypermethylated DMRs being associated with transcriptional increase, respectively (Supplemental_Table_S5.pdf). These DMRs are mainly of the CHGonly type (39/49 cases, 80%) and of the CGonly type (8/49 cases, 16%) (Supplemental_Table_S6.pdf). DMRs located upstream genes are mostly associated with transcriptional increase, but this is independent of the methylation change observed, with similar proportions of transcription increase for hypomethylated cases (67%) and hypermethylated cases (75%) (Supplemental_Table_S5.pdf). In contrast, DMRs located downstream genes show a co-occurrence of signal with transcription: 72% of hypermethylated DMRs co-occur with transcriptional increase, and 67% of hypomethylated DMR co-occur with transcriptional decrease (Supplemental_Table_S5.pdf). DMRs located upstream or downstream genes are mainly of the CG+CHG type, suggesting that the impact of this methylation change differs upon its location towards gene, with a more important impact when located downstream genes.

## DISCUSSION

Changes of DNA methylation following plant exposure to abiotic constraint has been observed for various species, different constraints and genomic features such as genes and TEs. However, these studies were either focusing on specific analysis of particular genes or TEs, or on detecting local DMRs from genome-scale datasets, therefore hampering the genome-wide analysis of the methylome response. Here, instead of relying on a window-based approach that does not reflect the feature-based biological role of the epigenome (Roudier et al. 2011), we took advantage of the genomic architecture of the maize genome to unravel how genomic features and chromosomal location may affect methylome response to cold. For the first time, we show that an abiotic stress induces a genome-wide hypermethylation through the hypermethylation of TE sequences across all chromosomes. This extensive modification is likely due to the long duration of the constraint applied, allowing for occurrence of cellular divisions under stress, and thus for the methylation pattern to amplify by increased methylation maintenance through each DNA replication. Use of an abiotic constraint inducing a clear physiological stress may also participate in this global response.

Our work highlights a specific response of TEs depending on their chromosomal location, with heterochromatic TEs undergoing hypermethylation at CGs and CHGs and TEs of the genic regions undergoing hypermethylation at CHHs. Mutant-based studies in standard conditions have shown that TE methylation is mostly dependent on local chromatin type (Sigman and Slotkin 2016). Typically, RdDM-dependent DNA methylation targets TEs located within or near genes (Gent et al. 2014), while methylation of heterochromatic TEs involves the action of methyltransferase MET1, chromomethylases CMT2 and CMT3, and chromatin remodeler DDM1 (Stroud et al. 2013; Zemach et al. 2013; Gent et al. 2014). Characteristics of the methylation increase that we observe following cold exposure supports this model, and provides new insights on the parallel reinforcement of both RdDM and non-RdDM methylation maintenance in response to prolonged cold in maize, leading to hypermethylation of TEs in all the chromatin regions where they reside.

TEs located inside genes are typically methylated in the three contexts with patterns similar to these located near genes, suggesting similar regulation (To et al. 2015; Sigman and Slotkin 2016). Here, we show that these two categories of TEs exhibit different methylation response to prolonged cold, pinpointing a difference in regulation. This difference is visible at CHGs, and particularly striking at CHHs, which methylation is almost unchanged for genic TEs, while near-gene TEs show methylation increase at the so-called CHH islands. These have been previously associated with DNA transposons and “spreading” LTR retrotransposon in maize (Li et al. 2015), which are enriched in the RLX type (Eichten et al. 2012). CHH island distance to gene was also shown to vary with TE indels among genotypes (Li et al. 2015). Here, we show that DNA methylation at CHH islands is increased by low temperature at most DNA transposons and RLX LTR retrotransposons, and that the level of the methylation change depends on the TE-gene distance. Our results therefore confirm that the strength of the RdDM regulation differs among TE types, but, most importantly, that the strength of the RdDM-based euchromatin-to-heterochromatin boundary depends on the TE-gene distance itself. This suggests that any change in the TE-gene distance (through the insertion or deletion of any type of sequence) likely influences the methylation levels at CHH islands. Interestingly, we did not observe a shift of CHH position in stressed plants, but rather an increase in DNA methylation at CHH island original position. This points to a reinforcement of RdDM at TE sequences themselves rather than modifying euchromatin-to-heterochromatin boundaries definition. Increase of methylation at CHH islands occurred in all categories of gene expression variation and did not reflect the transcriptional changes observed at neighboring genes (Supplemental_Fig_S13.pdf), thus confirming the lack of association between CHH islands and gene transcription levels, as observed from mutant analysis and within and between-species comparisons (Li et al. 2015; Niederhuth et al. 2016). This highlights the role of RdDM in regulating TE boundaries near genes, dependently to the distance to the closest gene but without an effect on the expression level of the gene.

Our results also provide first characterization of strong mCHG increase at all CENH3-rich regions described previously (Zhao et al., 2015). The underlying mechanism remains to be elucidated, but may rely on eviction or degradation of CENH3 and its replacement by H3. Depletion of CENH3 would allow for changing the overall chromatin architecture at these regions, leading to a better accessibility of H3 variants by CMT3 and/or helping the CHG/H3K9me2 reinforcing loop at H3 histones interacting with CENH3 at local level. Indeed, only a few hundreds of CENH3 occupy the centromere (Gent et al. 2012), allowing for large response of the H3 variants. In addition, because CENH3 histone tail does not contain H3K9, its replacement by H3 could potentially increase the mCHG reinforcing loop mediated by H3K9me2. Further studies will reveal whether CENH3-rich regions are sensitive to other abiotic stresses.

Together with these genome-wide changes, we provide evidence for local methylation remodeling, with characterization of 699 short regions (DMRs) showing differential methylation. As for landscapes variation, the vast majority of these DMRs are associated to TEs. The most abundant cases (57%) vary in all three contexts, and are likely traces of RdDM-spreading from TE edges to flanking sequences (Stroud et al. 2013; Zemach et al. 2013). Another 35% showing a mCHG change in high mCG regions, are associated with genic TEs, and likely occur from mis-regulation of the H3K9me2/IBM1 edge (To et al. 2015; Sigman and Slotkin 2016) leading to TE methylation spreading and epiallele formation. In contrast, CG only DMRs occur within genes in regions unmethylated at non-CG sites, and are in part associated with presence of a genic TE. This sustains the idea that CG gene-body methylation possibly evolved from silencing of ancient TE insertions (Bewick and Schmitz 2017; Niederhuth and Schmitz 2017), and highlights that response to abiotic stress may participate in maintaining gene body methylation in plants. Observation of similar amounts of hyper-and hypo-methylated DMRs highlights that methylation boundaries are mis-regulated but not oriented towards larger TE spreading. The two types of DMRs have been observed in other studies, for instance in *Arabidopsis* reponse to drought (8 hyper-and 9 hypo-methylated cases for CG or CHG DMRs, Ganguly et al. 2017) and rice response to phosphate deficit (147 hyper-vs. 28 hypo-methylated cases, Secco et al. 2015). In rice, hypermethylated DMRs were associated to TEs but not hypomethylated ones (Secco et al. 2015), suggesting different underlying mechanisms. In apple, drought induced similar amounts of hypo-and hyper-methylated DMRs in genes, but a larger amount of hypomethylated DMRs in TEs (Xu et al. 2018). This contrasts with our observation that hypo-and hypermethylated DMRs share the same annotation and cytosine contexts patterns, and are therefore likely generated from similar mechanisms. This suggests that generation of local methylation changes in response to abiotic constraints may be more complex than previously anticipated. Further analyses of these previously acquired datasets using the same methodology should help deciphering to what extent TE methylation spreading is affected by abiotic stresses in various species.

The methylome change we depict occurs together with extensive expression modifications. Interestingly, low temperature induces a transcriptional change for one third of the genes involved in the regulation of DNA methylation. These changes mainly correspond to an increase in expression, and affect genes involved in the RdDM pathway as well as in non-RdDM CG and CHG methylation maintenance. Expression change of genes involved in DNA methylation pathways has been reported for other abiotic constraints. For example, high temperature induces a coordinated up-regulation of RdDM genes such as *Drm2, NRPD1* and *NRPE1* in *Arabidopsis* (Naydenov et al. 2015), and osmotic stress reduces the expression of *Met1* and *Cmt3* orthologs in rice roots and shoots (Ahmad et al. 2014). In maize, four of the expressed methyltransferases (*ZmMet1a, ZmMet1b, ZmMet2a* and *ZmMet2b*) showed decreased expression levels in leaves after drought and salt stress (Qian et al. 2014). In our study, the transcriptional change affects a larger set of genes, involved in a wider set of methylation pathways, suggesting that prolonged cold has a more extensive effect on the expression of the methylation machinery. This has also been observed in apple grown under drought stress (Xu et al. 2018). Interestingly, while in other maize studies, genes from paralogous pairs showed similar transcriptional pattern (Qian et al. 2014), in our study, we observe a cold-induced transcriptional change for only one paralog of the pairs encoding the MET1, CMT3 and DDM1 function. For CMT3, and DDM1 functions, the most activated copy in our study was previously reported as the most highly transcribed (Li et al. 2014). For the DRM function, we observe a more complex response: *Zmet7/Dmt107/ZmDRM2/ZmMet3a*, which has been shown to be the most expressed (Li et al. 2014) remains unchanged in our experiment, while the *ZMet3/Dmt103/ZmDRM1/ZmMet3b* paralog shows a significant expression decrease and *ZMet6/Dmt106/ZmDRML/ZmMet3c* a significant expression increase. Similarly, genes encoding the IBM1 function show contrasted response: a significant expression increase for *Jmj22/Em_IBM1a* and significant expression decrease for *Jmj5/ZmIBM1b*. These results highlight the specificities of gene pairs involved in the regulation of methylation in maize following an abiotic stress, thus reinforcing the idea that gene duplication plays a role in maize adaptation.

Our work shows that prolonged low temperature induces extensive transcriptional activation of genes involved in the regulation of DNA methylation. This completely fits the methylome observations and highlights the parallel boost of these two pathways to methylate TEs in all the genomic locations where they reside. This transcriptional activation, together with the extensive methylation change at TEs genome-wide, shows the importance of hypermethylating TEs in response to an abiotic stress in a TE-dense genome as this of maize. Stress-induced transposition is a well-known phenomenon, and has been proposed to facilitate adaptation of populations to changing conditions (McClintock 1984). In a TE-dense genome, destabilization of a large number of TEs may nevertheless be too deleterious, which may explain why we observe a global hypermethylation of TEs. Interestingly, some TEs show methylation decrease, highlighting possible reactivation of few TEs while keeping the others hypermethylated. Destabilization of the *de novo* RdDM pathway at the transcriptional level is also observed in our experiment, including this of RNA polymerase II-related genes, and may also participate in reactivating TEs, as observed in *Arabidopsis* RNA polymerase II (Thieme et al. 2017). Further analysis of TE transcription will help decipher whether some TEs are transcriptionally reactivated in our experiment.

Whether the global increase in DNA methylation is involved in the phenotypic response of maize to low temperature remains to be fully elucidated. It may involve a molecular cost of genome-wide hypermethylation. Alternately, the local modification of methylation near genes involved in DNA repair, cell division control and cell wall composition may participate in the phenotypic response observed. Analyzing the phenotypic response of methylation mutants impaired in ZmMET1b, KYPb or MOP1 in our conditions may help answering this question. Comparing the response of genotypes with different levels of tolerance to cold treatment could also be very informative.

## METHODS

### Plant material and growth conditions

Seeds of maize inbred line B73 were provided by INRA, Saint-Martin-de-Hinx, France. Plants were grown in greenhouse (16h day at 24°C; 8h night at 18°C), for 4 days on petri dishes followed by 3 days in pots, until emergence stage (1-2 visible leaves). Then, plants for the stressed set were placed in an air conditioned room (16h day at 14°C; 8h night at 10°C) for a prolonged cold treatment of 22 days. Due to delay in development of the cold-treated plants, the unstressed set was sown with a shift of 18 days from the stressed set, to allow plants to reach similar developmental stages at sampling (Figure 1A). At the end of cold treatment, for both stressed and unstressed sets, samples were harvested independently from 3 distinct seedlings, directly frozen in liquid nitrogen and stored at −80°C for further analysis. Last visible leaf and one before last visible leaf samples were used for methylome and transcriptome analyses, respectively (Figure 1B).

### DNA extraction, and BS-seq data

DNAs were extracted from the last visible leaf samples following manufacturer’s protocol using Macherey Nagel midi extraction kit. For each sample, 5 µg of genomic DNA was fragmented by acoustic shearing to a target value of 250 bp using a Covaris S220 (Covaris, Woburn, MA), according to manufacturer’s recommendations. Fragmented DNA was purified using the QIAQuick PCR purification kit (Qiagen, Valencia, CA). Then, DNA fragments underwent an NGS library preparation procedure consisting in end repair and methylated adaptor ligation using the TruSeq DNA Sample Preparation kit (Illumina, San Diego, CA). Adapter-ligated fragments ranging from 200 to 300 nucleotides were isolated by agarose gel electrophoresis (Bio-Rad, Hercules, CA) and purified using the MinElute Gel Extraction Kit (Qiagen, Valencia, CA). Quality of the resulting NGS libraries was assessed by capillary electrophoresis using a Bioanalyzer DNA High-Sensitivity Chip (Agilent, Santa Clara, CA) and quantified using the Quant-It PicoGreen (Life Technologies, Carlsbad, CA). 500 ng of each native library were treated by sodium bisulfite using the EpiTect Bisulfite kit (Qiagen, Valencia, CA) and purified according to manufacturer’s instructions, followed by an additional round of purification using the MinElute PCR Purification Kit with a final resuspension volume of 15 µl (Qiagen, Valencia, CA). DNA fragments carrying both adapters after sodium bisulfite conversion were enriched by 8 cycles of PCR using the Pfu Turbo Cx Hotstart DNA Polymerase (Agilent, Santa Clara, CA). Primer dimers were subsequently removed using two rounds of purification using AMPure beads (Beckman Coulter, Indianapolis, IN). Quality and quantity of this amplified library was assessed as above. NGS data were generated by Illumina paired-end sequencing (2×100 bases) on a HiSeq2000 machine. Sequence quality was evaluated using FastQC (http://www.bioinformatics.babraham.ac.uk/projects/fastqc).

### BS-seq reads mapping and extraction of methylation levels

We generated between 226,087,680 and 497,596,575 read pairs (2 x 100 bases) per sample, which were mapped onto the B73 genome sequence (AGPv2). Fasta file from this genome sequence was indexed with the novoindex tool (http://www.novocraft.com/documentation/novoindex/) using the bisulfite option. Bisulfite treated reads were then aligned onto these two indexed references using Novoalign v3.04.06 using plant adapted stringent parameters. Alignment resulting files were split according to the mapping strand (CT or GA) and converted into bam compressed files using Samtools 1.2 (Li et al. 2009). Uniquely mapped reads were kept for further analysis. Samtools flagstat (http://www.htslib.org/doc/samtools.html) was used to assess the quantity of paired, properly paired or single reads among these uniquely mapped reads. Methylation level per cytosine was estimated by Novomethyl following the Novocraft protocol (http://www.novocraft.com/documentation/novoalign-2/novoalign-user-guide/bisulphite-treated-reads/novomethyl-analyzing-methylation-status/). An in-house perl script was then used to exclude heterozygous sites and low quality consensus calling. Bisulfite conversion rate was above 99.7% in all samples, as estimated from chloroplast cytosine analysis. Finally, methylation files from the 6 samples were split by cytosine context, and cytosines with 4x to 25x coverage depth in all 6 samples were extracted using a custom perl script.

### Profiling methylation difference between stressed and unstressed plants

For comparison of genome-wide methylation levels, we performed two analyses: (i) we computed average methylation levels for each sample from all cytosines of the genome, using each context independently and (ii) we compared methylated cytosine numbers using a Student’s test, considering cytosines with methylation levels above 80% for CG, 70% for CHG and 5% for CHH as methylated. For chromosome landscapes, average methylation levels per cytosine were calculated for the 3 stressed and for the 3 unstressed samples, and the difference between average values of stressed and unstressed samples was computed for 100kb windows (genome) or over genomic features size. Features with less than 1 CHH per 100bp were eliminated from the analysis. TE and gene positions were extracted from B73 AGPv2 annotation (http://ftp.maizesequence.org/release-5b/). Gene cytosines overlapping with TEs were extracted for each context using Bedtools intersect (Quinlan and Hall 2010) and removed from the gene set. Genomic regions with no overlap with gene or TE annotations were extracted and considered as “other” type of feature. Extraction of chromosomic regions with outlying CHG variation was performed by segmentation analysis using the R DNAcopy package (https://bioconductor.org/biocLite.R). These segments were used to compare methylation variation between centromeric outlying segments and the rest of the genome for the different underlying features. CHH methylation patterns in genes and 2kb flanking regions were computed from average methylation values of stressed and unstressed plants using Deeptools (Ramirez et al. 2014) after classification of genes based on their expression variation (see below). Genes were oriented from TSS to TTS using the strand information from the corresponding .gff annotation file.

### Detection of Differentially Methylated Regions (DMRs) between “stressed” and “unstressed” plants

Differentially Methylated Regions (DMRs) and Differentially Methylated Cytosines (DMCs) were detected between “stressed” and “unstressed” plants with the methylKit R package (Akalin et al. 2012). CG and CHG cytosines were used separately, and a q-value below 0.01 and a methylation difference over 10% were required. For DMRs, sliding windows of 200, 500 and 1000 bp with a step size of 100 bp were used. Number of DMCs per DMR was estimated by comparing DMRs and DMCs positions using Bedtools intersect (Quinlan and Hall 2010) with the option -c. DMRs with at least 5 DMCs were kept for further analysis. Finally, overlapping DMRs detected in different window size experiments were merged together using a custom Python script. In this step, when the same amount of DMC was found in the three window sets, the DMR with shortest size was kept. Otherwise, the DMR with largest number of DMCs was kept. Methylation levels of these regions were then estimated for each sample with roimethstat (http://smithlabresearch.org/software/methpipe) using the original methylation call files of the three contexts.

### DMR classification by context

DMR classification into “CG only”, “CHG only” and “CG plus CHG” categories was performed by a 2-component Gaussian mixture model using the mixture R package. The Gaussian covariance structure was selected based on Bayesian Information Criterion (BIC). Each DMR was assigned to a cluster when its posterior probability was higher than 90%. It was considered unclassified otherwise. For “CG plus CHG” cases, redundancy between cases originally found with CG and originally found with CHG were merged using a 50% overlap threshold using Bedtools intersect (Quinlan and Hall 2010), keeping the most extreme boundaries. DNA methylation levels were re-estimated for these new locations using roimethstat (http://smithlabresearch.org/software/methpipe).

### DMR annotation

DMR location was compared to this of genes and transposons from B73 AGPv2 using Bedtools *intersect* (Quinlan and Hall 2010). DMRs were categorized into “inside”, “upstream”, “downstream” or “away” based on distance, strand and relative start and end positions using an in-house perl script. Threshold distance was 2 kb for genes and 500 bp for TEs. Enrichment of DMRs in the different categories was based on relative abundance as compared to a random set of regions. For each DMR, a random region with same size and from same chromosome was retrieved from the B73 genome using Bedtools *shuffle* (V.2.7.3) (Quinlan and Hall 2010) and subsequently checked to contain at least 5 cytosines from the [4x-25x] dataset used for original DMR detection using a custom Python script (5 GCs for “CG only” cases, 5 CHGs for CHG only cases, and 5 CGs and 5 CHGs for CG plus CHG cases). For analysis of DMRs located inside genes, new random sets were generated, excluding genomic regions located outside of genes.

### mRNA extraction, RNA-seq library preparation and RNA sequencing

mRNAs were extracted from the one before last visible leaf samples with Trizol (Invitrogen ref.15596018) and β-mercaptoethanol (SIGMA ref. M3148-25ML) reagents. Supernatant was recovered and RNA purified using Qiagen RNeasy Plant Mini kit (ref. 74904) following manufacturer’s instructions. Then, a Qiagen RNAse-free DNAse set (ref. 79254) was applied to remove the residual DNA. Library construction and sequencing were generated by the IPS2-POPS platform with Illumina NexSeq500 sequencer using barcoded adaptors and 18 samples per lane. Briefly, mRNA were polyA selected, fragmented to 260 bases and libraries were performed using the TruSeq stranded mRNA kit (Illumina®, California, U.S.A.) and sequenced in paired-end (PE) with 150 bases read length.

### mRNA_seq reads quality trimming and mapping

mRNA reads were trimmed for library adapters and for Phred Quality Score Qscore >20 and read length >30 bases using Trimmomatic version 0.36 (Bolger et al. 2014). Ribosome sequences were removed using sortMeRNA version 2.1 (Kopylova et al. 2012). Paired-ends reads were then aligned onto the B73 AGPv2 reference sequence. STAR version 2.5.2b was used for mapping, with options -- outSAMprimaryFlag AllBestScore --outFilterMultimapScoreRange 0 to keep the bests results. Only unambiguously mapped reads were used, and multihits were removed.

### Detection of differentially expressed genes

Differential analysis followed the procedure described in (Rigaill et al. 2018). Briefly, after counts per million (CPM) normalization was performed, genes with less than one read in at least one half of the samples were discarded. Library size was normalized using the trimmed mean of M-value (TMM) method and count distribution was modeled with a negative binomial generalized linear model where the environment factor (“stressed” or “unstressed” plants) was taken into account. Dispersion was estimated by the edgeR method (Version 1.12.0, (McCarthy et al. 2012) in the ‘R’ statistical software (http://www.R-project.org/). Expression differences were compared between stressed and unstressed plants using likelihood ratio test and p-values were adjusted by the Benjamini-Hochberg procedure to control False Discovery Rate (FDR). Genes with adjusted p-value < 0.05 were declared differentially expressed. Genes were then subdivided into “expressed without differential expression”, “with up differential expression” and “with down differential expression” to perform CHH islands analyses (see above). A list of 80 genes (Supplemental_Table_S4.pdf) described in the literature as involved in DNA methylation regulation in maize (Singh et al. 2011; Candaele et al. 2014; Haag et al. 2014; Li et al. 2014; Petsch et al. 2015) or extracted from maizeGDB (https://www.maizegdb.org/) by searching for names of classical genes involved in Arabidopsis DNA methylation pathways (Law and Jacobsen 2010; Sigman and Slotkin 2016), and compared to the list of differentially expressed genes. Counts presented in Table 2 were also normalized by gene length to get comparable estimate of gene expression levels among genes in stressed and unstressed plants.

### Gene ontology analysis

Enrichment tests in gene ontology categories were performed with TopGo and Rgraphviz R packages (http://bioconductor.org/biocLite.R), using either the list of up differentially expressed genes, down differentially expressed genes, genes with a hypermethylated DMR within 2kb and genes with a hypomethylated DMR within 2kb separately. Gene ontology information was retrieved from http://ftp.maizesequence.org/release-5b/.

## DATA ACCESS

All raw sequencing data (bisulfite-seq and mRNA-seq) generated in this study are available on request.

## ACKNOWLEDGEMENTS

We thank INRA Saint Martin de Hinx for seed stocks. The POPS platform benefits from the support of the LabEx Saclay Plant Sciences-SPS (ANR-10-LABX-0040-SPS). We thank Daniel Grimanelli for providing a list of DNA methylation genes, Kelly Dawe for discussion on centromeres, and Karine Alix, Nicolas Bouché and Leandro Quadrana for helpful discussions on the manuscript.

