## Supplemental_Tables_and_Figures for "Low temperature triggers genome-wide hypermethylation of transposable elements and centromeres in maize"

**Achour et al. Supplemental Material**

**Table of contents:**

**Supplemental Table S1 : Global methylation in the 6 samples**

**Supplemental Table S2: DMR characteristics**

**Supplemental Table S3: DMR classification**

**Supplemental Table S4: List of 80 genes involved in regulation of DNA methylation in maize**

**Supplemental Table S5: Comparison of DMRs location and gene expression change**

**Supplemental Table S6: Comparison of DMRs location and gene expression change, classified by CGonly, CHGonly and CG+CHG**

**Supplemental Figure S1: Methylation landscape of chromosomes 1 to 10**

**Supplemental_Figure_S2: Chromosome landscapes of average methylation difference for CG, CHG and CHH cytosine contexts and for the different types of genomic features**

**Supplemental Figure S3: Distribution of methylation difference in TEs for CG, CHG and CHH contexts with respect to distance to closest gene**

**Supplemental Figure S4: Methylation difference for hypermethylated TEs with respect to gene location and cytosine context**

**Supplemental Figure S5: Methylation difference for hypomethylated TEs with respect to gene location and cytosine context**

**Supplemental Figure S6: Distribution of DMR size in hyper- (red) and hypomethylated (blue) DMR cases**

**Supplemental Figure S7: Distribution of average methylation difference hyper- (red) and hypomethylated (blue) DMR cases**

**Supplemental Figure S8: Distribution of cytosine number in hyper- (red) and hypomethylated (blue) DMR cases**

**Supplemental Figure S9: Distribution of DMC number in hyper (red) and hypo (blue) DMR cases**

**Supplemental Figure S10: Comparison of average CG and CHG methylation difference between stressed and unstressed plants for hypermethylated DMR cases**

**Supplemental Figure S11: Comparison of average CG and CHG methylation difference between stressed and unstressed plants for hypomethylated DMR cases**

**Supplemental Figure S12: Characteristics of hypomethylated DMRs**

**Supplemental Figure S13: Average CHH methylation levels within genes and in flanking regions**

**<u>Supplemental Table S1</u> : Global methylation in the 6 samples**

U: Unstressed plants, S: Stressed plants

| Sample | Number of cytosines analyzed | | | Genome-wide average methylation level | | | Number of methylated cytosines | | | Percent of methylated cytosines | | | Fisher's test p-value | | | Student's test p-value | | |
|---|---|---|---|---|---|---|---|---|---|---|---|---|---|---|---|---|---|---|
| | CG | CHG | CHH | CG | CHG | CHH | CG | CHG | CHH | CG | CHG | CHH | CG | CHG | CHH | CG | CHG | CHH |
| B73_U_rep1 | 54 444 473 | 44 451 905 | 148 406 979 | 82.08 | 68.98 | 1.17 | 44 688 548 | 29 258 868 | 10 542 362 | 82.08 | 65.82 | 7.10 | 0.31 | 0.74 | 0.01 | 0.19 | 0.03 | 0.15 |
| B73_U_rep2 | | | | 80.93 | 67,00 | 0.98 | 43 018 217 | 27 957 895 | 9 489 133 | 79.01 | 62.89 | 6.39 | | | | | | |
| B73_U_rep3 | | | | 80.78 | 67.23 | 0.98 | 42 620 101 | 28 137 326 | 9 790 534 | 78.28 | 63.30 | 6.60 | | | | | | |
| B73_S_rep1 | | | | 82.79 | 71.01 | 1.2 | 44 700 947 | 30 349 943 | 10 682 882 | 82.10 | 68.28 | 7.20 | | | | | | |
| B73_S_rep2 | | | | 82.94 | 71.26 | 1.19 | 44 919 371 | 30 479 139 | 10 673 699 | 82.50 | 68.57 | 7.19 | | | | | | |
| B73_S_rep3 | | | | 81.5 | 69.35 | 1.09 | 44 012 376 | 29 482 710 | 10 621 782 | 80.84 | 66.32 | 7.16 | | | | | | |

**<u>Supplemental Table S2:</u> DMR characteristics**

DMR: Differentially Methylated Region, DMC: Differentially methylated Cytosine

| Type | Hypermethylated | | Hypomethylated | |
|---|---|---|---|---|
| Context | CG | CHG | CG | CHG |
| DMR number | 192 | 162 | 228 | 168 |
| Median DMR size (bp) | 400 | 800 | 400 | 700 |
| Median methylation difference (%) | 29 | 35 | 25 | 30 |
| Median cytosine number per DMR | 21 | 34 | 30 | 39 |
| Median DMC number | 6 | 7 | 7 | 7 |

**<u>Supplemental Table S3:</u> DMR classification**

| | Hypermethylated DMRs | | Hypomethylated DMRs | | All DMRs | |
|---|---|---|---|---|---|---|
| Type | Number | Percent total | Number | Percent total | Number | Percent total |
| CG only | 30 | 9.3 | 18 | 4.8 | 48 | 6.9 |
| CHG only | 100 | 31.0 | 120 | 31.8 | 220 | 31.5 |
| CG plus CHG | 180 | 55.9 | 183 | 48.5 | 363 | 51.9 |
| Unclassified | 12 | 3.7 | 56 | 14.8 | 68 | 9.7 |
| Total | 322 | | 377 | | 699 | |

**<u>Supplemental Table S4:</u> List of 80 genes involved in regulation of DNA methylation in maize**

| | Name: | | | | | |
|---|---|---|---|---|---|---|
| **Gene ID** | **in MaizeGDB** | **in Haag et al., 2014** | **in Li et al., 2014** | **in Candaele et al., 2014** | **in Qian et al., 2014** | **from D. Grimanelli pers. comm.** |
| GRMZM2G141818 | Ago104 | | | | | |
| GRMZM2G589579 | Ago4a | Ago105 | | | | |
| GRMZM2G089743 | Ago105 | Ago119 | | | | |
| GRMZM2G347402 | Ago6 | Ago121 | | | | |
| GRMZM5G574858 | Chr127 | Chr127/Zm_Drd1a | | | | |
| GRMZM2G393742 | Chr156 | Chr156/Zm_Drd1b | | | | |
| GRMZM2G178435 | Chr167 | Chr167/Rmr1-like | | | | |
| GRMZM2G040762 | Dcl101 | | | | | |
| GRMZM2G413853 | Dcl102 | | | | | |
| GRMZM2G160473 | Dcl4 | | | | | |
| GRMZM5G814985 | Dcl104 | | | | | |
| GRMZM2G301405 | Dcl105 | | | | | |
| GRMZM2G024466 | Dcl4 | | | | | |
| GRMZM2G050869 | Dcl4 | | | | | |
| GRMZM2G050882 | Dcl4 | | | | | |
| GRMZM2G177165 | Chr101 | | | | | Zm_DDM1a |
| GRMZM2G071025 | Chr106 | | | | | Zm_DDM1b |
| GRMZM2G309152 | no name | Zm_Dms3 | | | | |
| GRMZM2G417089 | Jmj22 | | | | | Zm_IBM1a |
| GRMZM2G070885 | Jmj5 | | | | | Zm_IBM1b |
| GRMZM2G098603 | no name | Iwr1/Zm_Dms4/Zm_Rdm4 | | | | |
| GRMZM2G054383 | no name | | | | | SUVH4/5/6/Zm_KYPa |
| GRMZM2G336909 | no name | | | | | SUVH4/5/6/Zm_KYPb |
| GRMZM2G333916 | Met1 | | | ZmMet2 | ZmMet1a | |
| GRMZM2G334041 | Met8 | | Zmet1 | ZmMet1 | ZmMet1b | |
| GRMZM2G025592 | Met2 | | Zmet2/Dmt102 | ZmCMT1 | ZmMet2a | |
| GRMZM2G005310 | Met5 | | Zmet5/Dmt105 | ZmCMT2 | ZmMet2b | |
| GRMZM2G092497 | Met3 | | Zmet3/Dmt103 | ZmDRM1 | ZmMet3b | |
| GRMZM2G137366 | Met7 | | Zmet7/Dmt107 | ZmDRM2 | ZmMet3a | |
| GRMZM2G065599 | Met6 | | Zmet6/Dmt106 | ZmDRML | ZmMet3c | |
| GRMZM2G157589 | Met4 | | | ZmDNMT2 | ZmMet4 | |
| GRMZM2G042443 | Mop1 | | | | | |
| NP_001152395 | no name | Nrp(b/d/e)10a | | | | |
| GRMZM2G130207 | no name | Nrp(b/d/e)11 | | | | |
| GRMZM2G402295 | no name | Nrp(b/d/e)3 | | | | |
| GRMZM2G013600 | no name | Nrp(b/d/e)6a | | | | |
| GRMZM2G086904 | no name | Nrp(b/d/e)6b | | | | |
| GRMZM2G034326 | no name | Nrp(b/d/e)8 | | | | |
| GRMZM5G803992 | Umc2069 | Nrp(d/e)10b | | | | |
| GRMZM5G834335 | no name | Nrp(d/e)10c | | | | |
| GRMZM2G054225 | Nrpd2/e2 | Nrp(d/e)2a/Mop2/Rmr7 | | | | |
| GRMZM2G427031 | no name | Nrp(d/e)2b | | | | |
| GRMZM2G453424 | no name | Nrp(d/e)4 | | | | |
| GRMZM2G469969 | gldh1 | Nrp(d/e)5 | | | | |
| GRMZM2G040702 | no name | Nrp(d/e)7 | | | | |
| GRMZM5G898768 | Ufg57 | Nrp(d/e)9 | | | | |
| GRMZM2G044306 | no name | Nrpb1 | | | | |
| GRMZM2G043461 | no name | Nrpb11-like | | | | |
| GRMZM2G146331 | no name | Rpb12-like | | | | |
| GRMZM2G322661 | no name | Rpb12-like | | | | |
| GRMZM2G540834 | no name | Rpb12-like | | | | |
| GRMZM2G084891 | no name | Nrpb2a | | | | |
| GRMZM2G113928 | no name | Nrpb2b | | | | |
| GRMZM2G119393 | no name | Nrbp4 | | | | |
| GRMZM2G476009 | no name | Nrpb5a | | | | |
| GRMZM2G099183 | no name | Nrpb5b | | | | |
| GRMZM2G179346 | no name | Nrpb7 | | | | |
| GRMZM2G347789 | no name | Nrpb8-like | | | | |
| GRMZM2G046061 | no name | Nrpb9a | | | | |
| GRMZM2G023028 | no name | Nrpb9b | | | | |
| GRMZM2G007681 | Rmr6 | Rmr6/Nrpd1 | Mop3 | | | |
| GRMZM2G153797 | no name | Nrpe1 | | | | |
| GRMZM2G133512 | no name | Nrpe2c | | | | |
| GRMZM2G145201 | Rdr6 | Rdr102 | | | | |
| GRMZM2G154946 | Rmr1 | Rmr1 | | | | |
| GRMZM2G131756 | Ros1 | | | | | |
| GRMZM2G047104 | no name | | | | | Zm_SHH1 |
| GRMZM2G111204 | Hb131 | Zm_SHH2a | | | | |
| GRMZM2G126170 | Hb55 | Zm_SHH2b | | | | |
| GRMZM5G843174 | Magi73349 | | | | | SHH2c |
| GRMZM2G025924 | Sdg137 | | | | | Zm_SUVH2/9 |
| GRMZM2G034288 | Sdg136 | | | | | Zm_SUVH2/9 |
| GRMZM2G021044 | Sdg104 | | | | | Zm_SUVH5/6 |
| GRMZM2G054380 | Sdg118 | | | | | Zm_SUVH5/6 |
| GRMZM2G074094 | Sdg103 | | | | | Zm_SUVH5/6 |
| GRMZM2G300955 | Sdg119 | | | | | Zm_SUVH5/6 |
| GRMZM2G162211 | no name | | | | | Zm_Vim1/2/3 |
| GRMZM2G339151 | Vim102 | | | | | |
| GRMZM2G461447 | Vim103 | | | | | |
| AC191534.3_FG003 | Vim104 | | | | | |

**<u>Supplemental Table S5:</u> Comparison of DMRs location and gene expression change**

DEGs: Differentially Expressed genes. nr: not relevant

| | Position of DMR as compared to gene | Number of DMRs associated with annotated genes | Number of DMRs associated with downregulated genes | Number of DMRs associated with upregulated genes | Number of DMRs associated with DEGs | DMRs associated with DEGs/DMRs associated with gene (%) | Upregulated genes/Total (%) |
|---|---|---|---|---|---|---|---|
| **Total DMRs (all)** | Inside | 223 | 26 | 49 | 75 | 33.6 | 65.3 |
| | Downstream | 81 | 15 | 14 | 29 | 35.8 | 48.3 |
| | Gene included in DMR | 3 | 0 | 0 | 0 | 0.0 | nr |
| | Upstream | 70 | 4 | 9 | 13 | 18.6 | 69.2 |
| | **Total** | **377** | **45** | **72** | **117** | **31.0** | **61.5** |
| **Hypomethylated DMRs (all)** | Inside | 111 | 15 | 26 | 41 | 36.9 | 63.4 |
| | Downstream | 36 | 12 | 6 | 18 | 50.0 | 33.3 |
| | Gene included in DMR | 1 | 0 | 0 | 0 | 0.0 | nr |
| | Upstream | 37 | 3 | 6 | 9 | 24.3 | 66.6 |
| | **Total** | **185** | **30** | **38** | **68** | **36.8** | **55.9** |
| **Hypermethylated DMRs (all)** | Inside | 112 | 11 | 23 | 34 | 30.4 | 67.6 |
| | Downstream | 45 | 3 | 8 | 11 | 24.4 | 72.7 |
| | Gene included in DMR | 2 | 0 | 0 | 0 | 0.0 | nr |
| | Upstream | 33 | 1 | 3 | 4 | 12.1 | 75.0 |
| | **Total** | **192** | **15** | **34** | **49** | **25.5** | **69.4** |

**<u>Supplemental Table S6:</u> Comparison of DMRs location and gene expression change, classified by CGonly, CHGonly and CG+CHG**

nr: not relevant

| | Position of DMR as compared to gene | Number of DMRs associated with annotated genes | Number of DMRs associated with downregulated genes | Number of DMRs associated with upregulated genes | Number of DMRs associated with DEGs | DMRs associated with DEGs/DMRs associated with gene (%) | Upregulated genes/Total (%) |
|---|---|---|---|---|---|---|---|
| **Hypomethylated DMRs (CHGonly)** | Inside | 78 | 12 | 20 | 32 | 41.0 | 25.6 |
| | Downstream | 6 | 1 | 0 | 1 | 16.7 | 0.0 |
| | Gene included in DMR | 0 | 0 | 0 | 0 | nr | nr |
| | Upstream | 4 | 0 | 0 | 0 | 0.0 | 0.0 |
| | **Total** | **88** | **13** | **20** | **33** | **37.5** | **22.7** |
| **Hypomethylated DMRs (CGonly)** | Inside | 12 | 0 | 4 | 4 | 33.3 | 33.3 |
| | Downstream | 1 | 1 | 0 | 1 | 100 | 0.0 |
| | Gene included in DMR | 0 | 0 | 0 | 0 | nr | nr |
| | Upstream | 1 | 0 | 0 | 0 | 0.0 | 0.0 |
| | **Total** | **14** | **1** | **4** | **5** | **35.7** | **28.6** |
| **Hypomethylated DMRs (CG+CHG)** | Inside | 21 | 3 | 2 | 5 | 23.8 | 9.5 |
| | Downstream | 29 | 10 | 6 | 16 | 55.2 | 20.7 |
| | Gene included in DMR | 1 | 0 | 0 | 0 | 0.0 | 0.0 |
| | Upstream | 32 | 3 | 6 | 9 | 28.1 | 18.8 |
| | **Total** | **83** | **16** | **14** | **30** | **36.1** | **16.9** |
| **Hypermethylated DMRs (CHGonly)** | Inside | 77 | 7 | 19 | 26 | 33.8 | 24.7 |
| | Downstream | 5 | 0 | 1 | 1 | 20.0 | 20.0 |
| | Gene included in DMR | 0 | 0 | 0 | 0 | nr | nr |
| | Upstream | 3 | 0 | 0 | 0 | 0.0 | 0.0 |
| | **Total** | **85** | **7** | **20** | **27** | **31.8** | **23.5** |
| **Hypermethylated DMRs (CGonly)** | Inside | 20 | 3 | 4 | 7 | 35.0 | 20.0 |
| | Downstream | 4 | 0 | 1 | 1 | 25.0 | 25.0 |
| | Gene included in DMR | 0 | 0 | 0 | 0 | nr | nr |
| | Upstream | 1 | 0 | 0 | 0 | 0.0 | 0.0 |
| | **Total** | **25** | **3** | **5** | **8** | **32.0** | **20.0** |
| **Hypermethylated DMRs (CG+CHG)** | Inside | 15 | 1 | 0 | 1 | 6.7 | 0.0 |
| | Downstream | 36 | 3 | 6 | 9 | 25.0 | 16.7 |
| | Gene included in DMR | 2 | 0 | 0 | 0 | 0.0 | 0.0 |
| | Upstream | 29 | 1 | 3 | 4 | 13.8 | 10.3 |
| | **Total** | **82** | **5** | **9** | **14** | **17.1** | **11.0** |

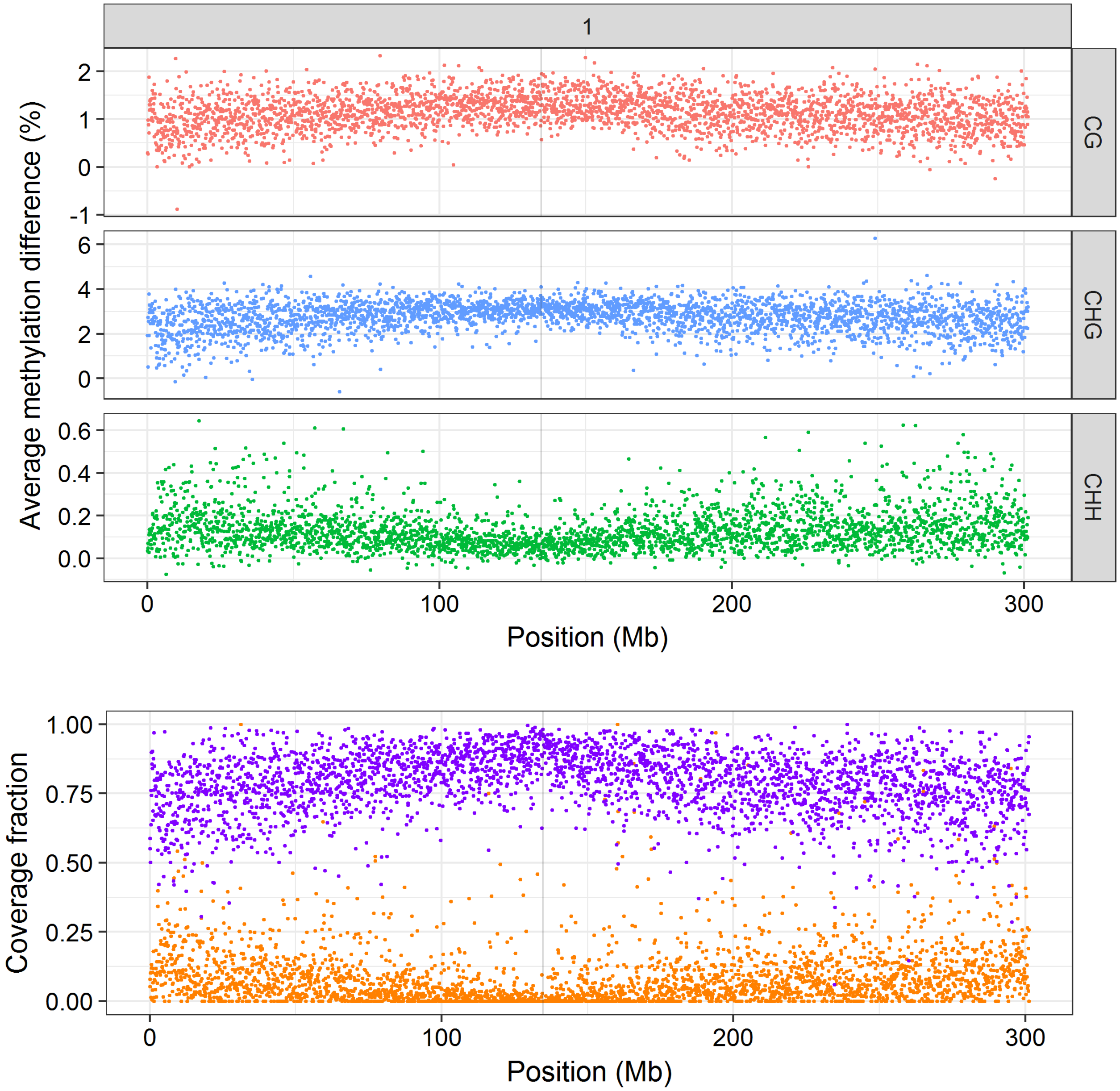


**<u>Supplemental Figure S1:</u> Methylation landscape of chromosomes 1 to 10**

Methylation landscapes showing average difference between methylation levels of stressed and unstressed plants over 100kb bins, for each of the three cytosine contexts, and annotation landscape showing percent of coverage of each 100kb bin by TE (purple) and gene (orange) bases. Contexts are shown on the right, in grey panels. Chromosome number is indicated on top. Positions are in Mb from tip of chromosomes. Centromere positions are highlighted by grey boxes.

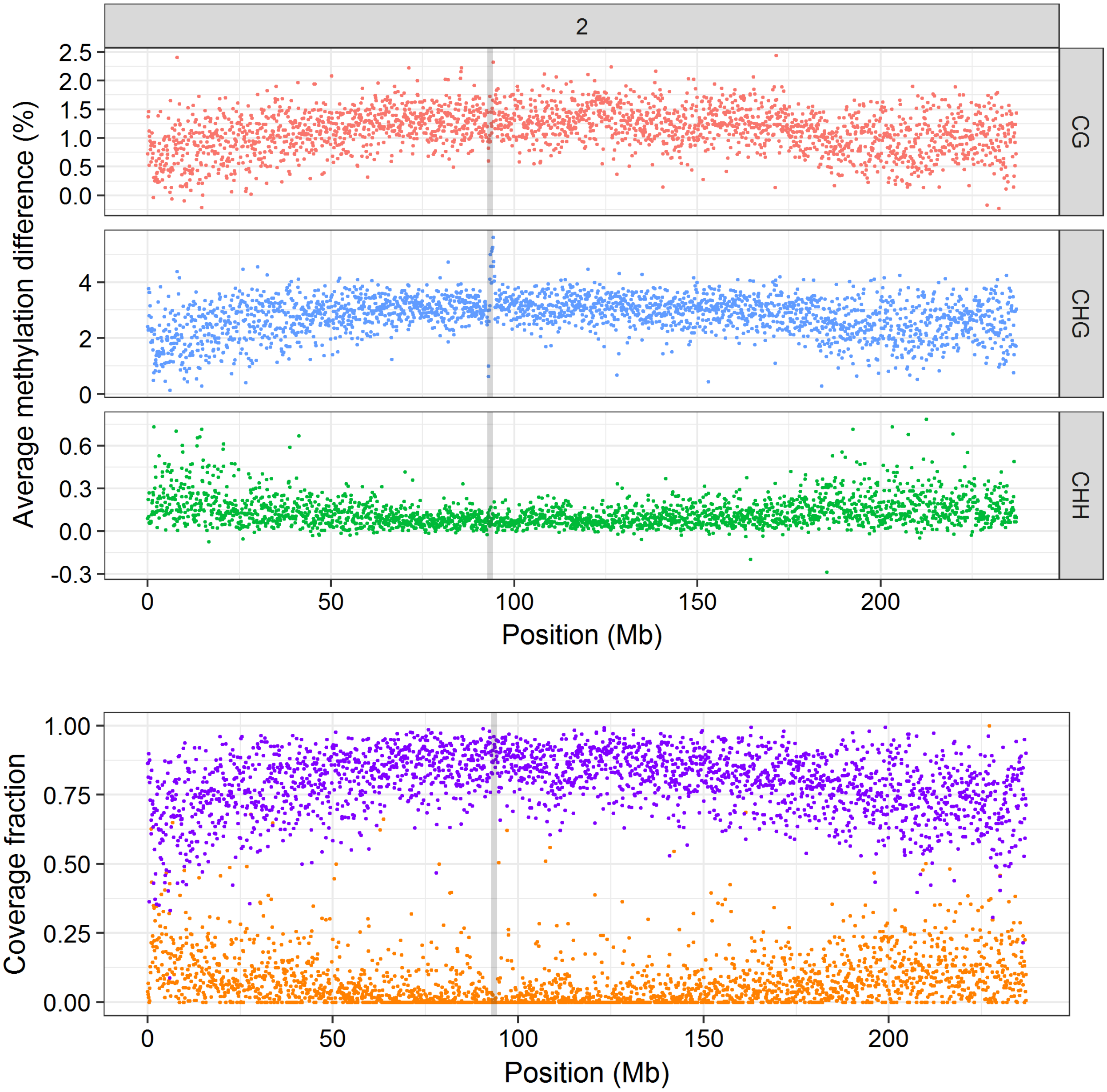


**<u>Supplemental Figure S1 (continued):</u>** **Methylation lanscape of chromosomes 1 to 10**

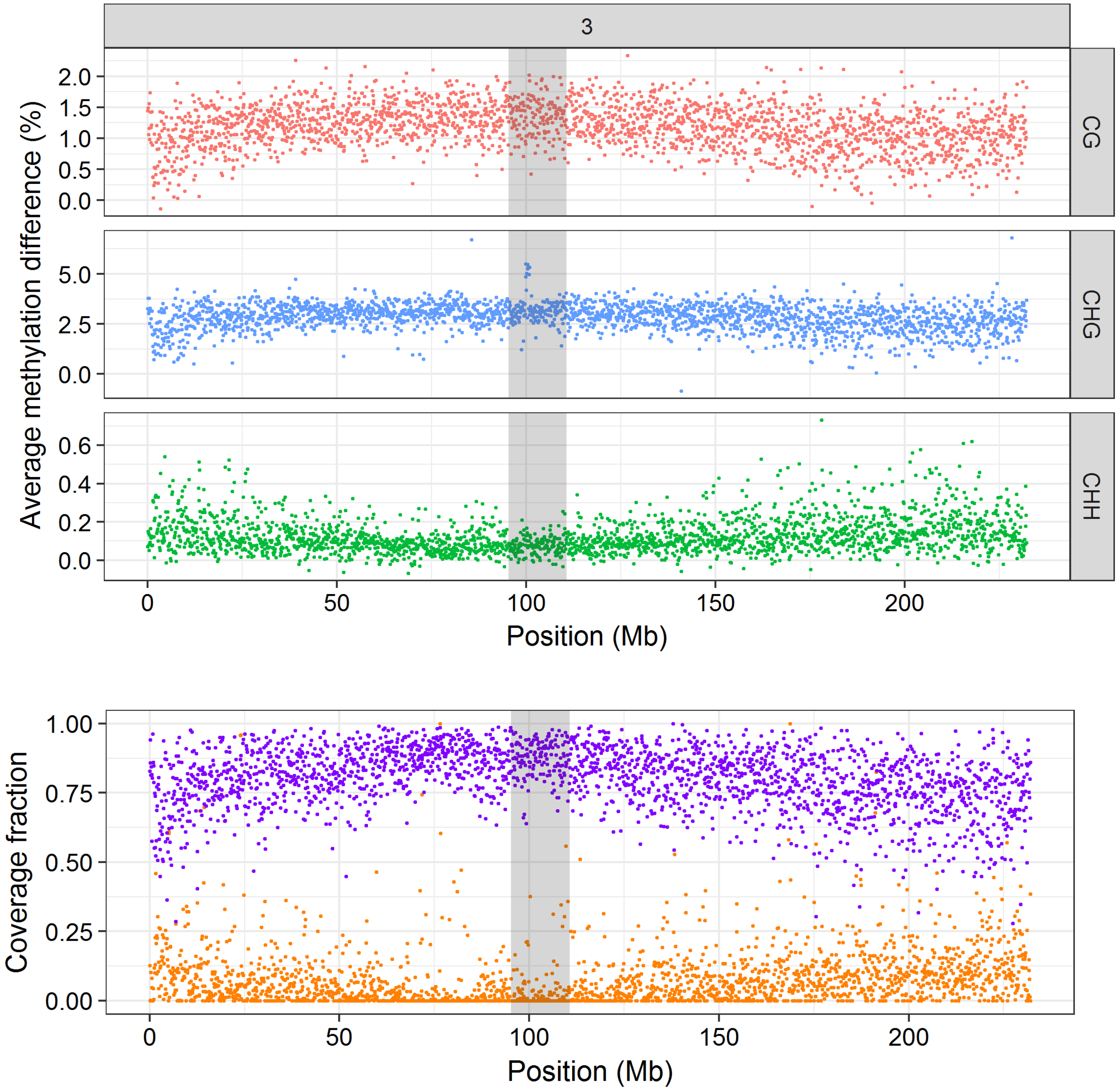


**<u>Supplemental Figure S1 (continued):</u>** **Methylation lanscape of chromosomes 1 to 10**

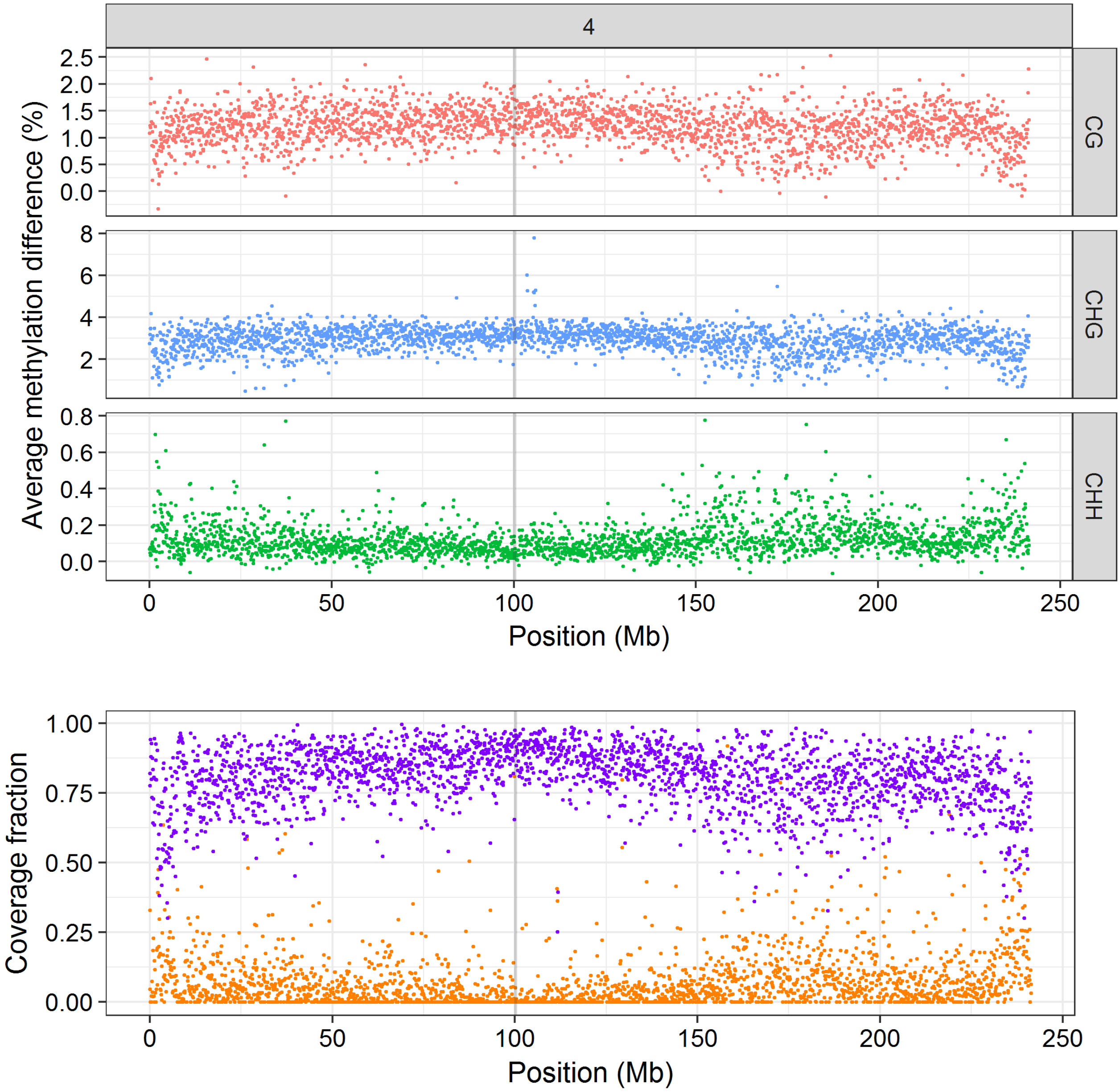


**<u>Supplemental Figure S1 (continued):</u> Methylation lanscape of chromosomes 1 to 10**

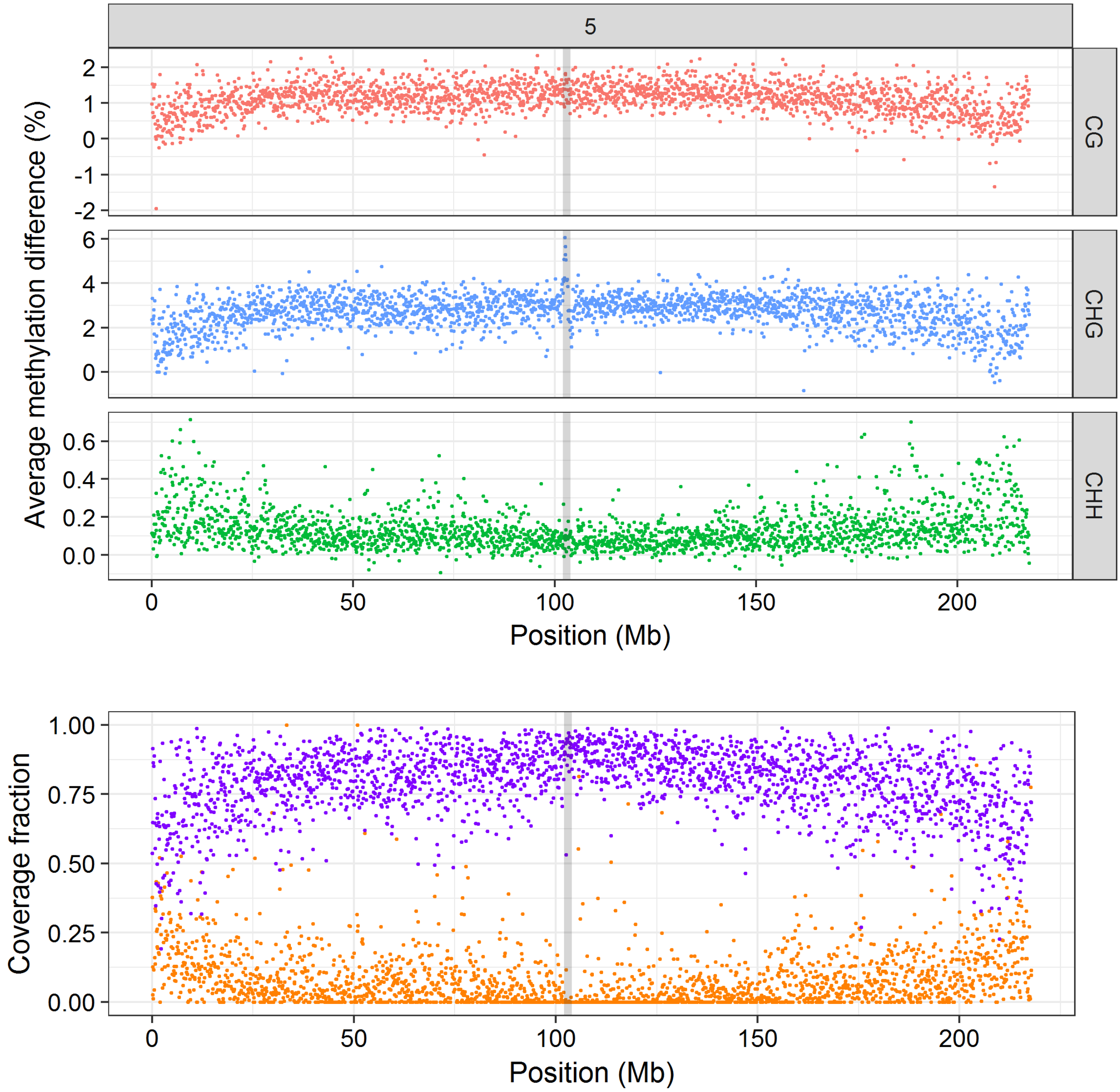


**<u>Supplemental Figure S1 (continued):</u>** **Methylation lanscape of chromosomes 1 to 10**

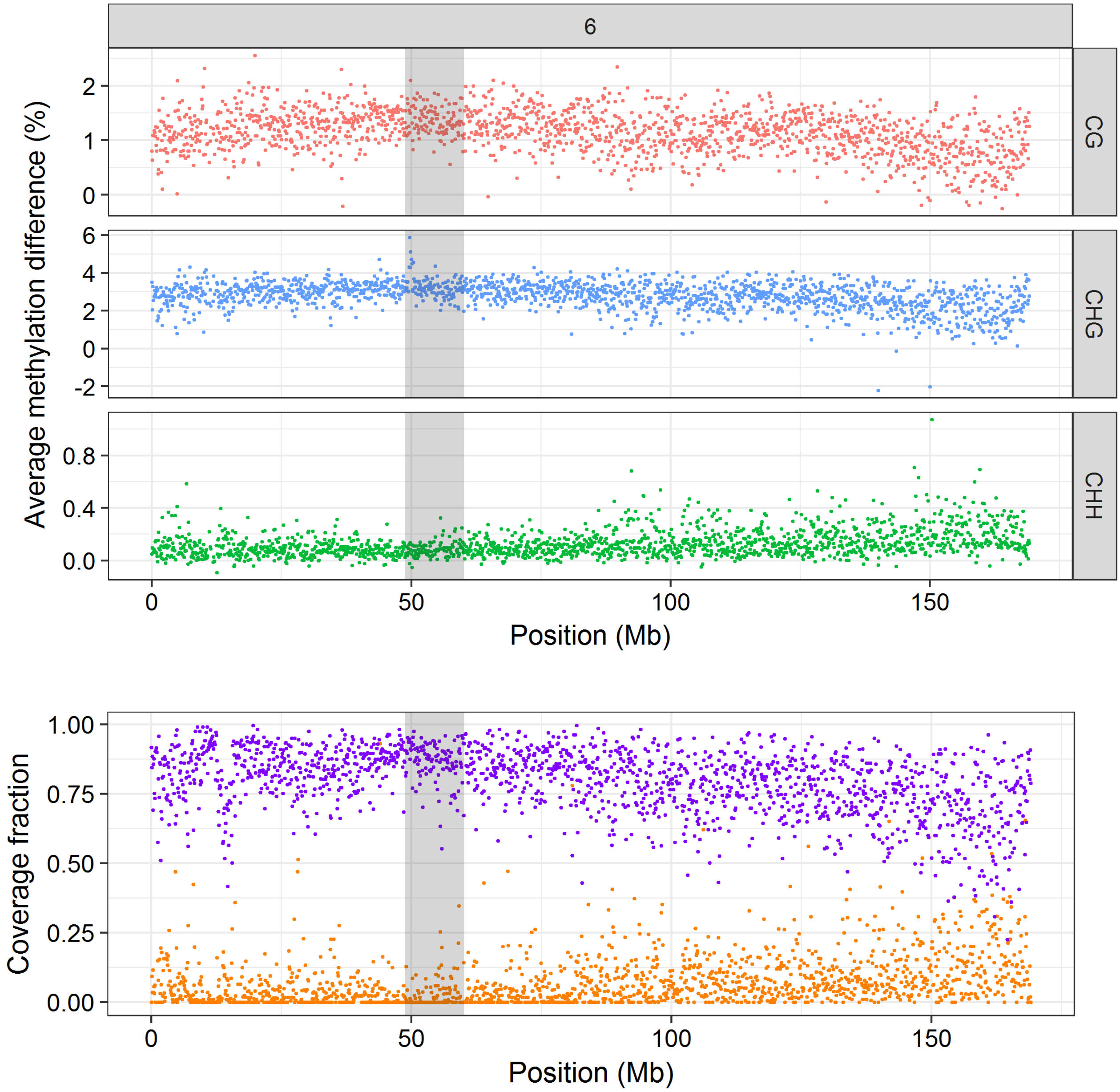


**<u>Supplemental Figure S1 (continued):</u>** **Methylation lanscape of chromosomes 1 to 10**

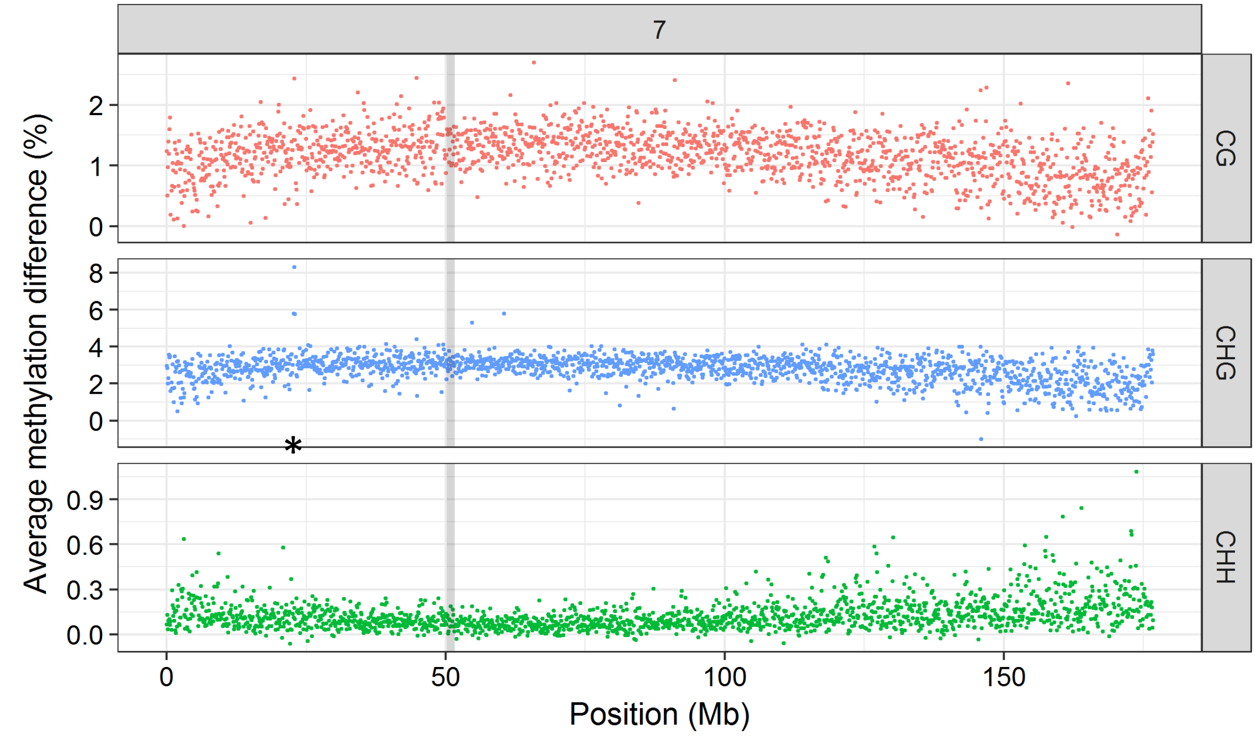


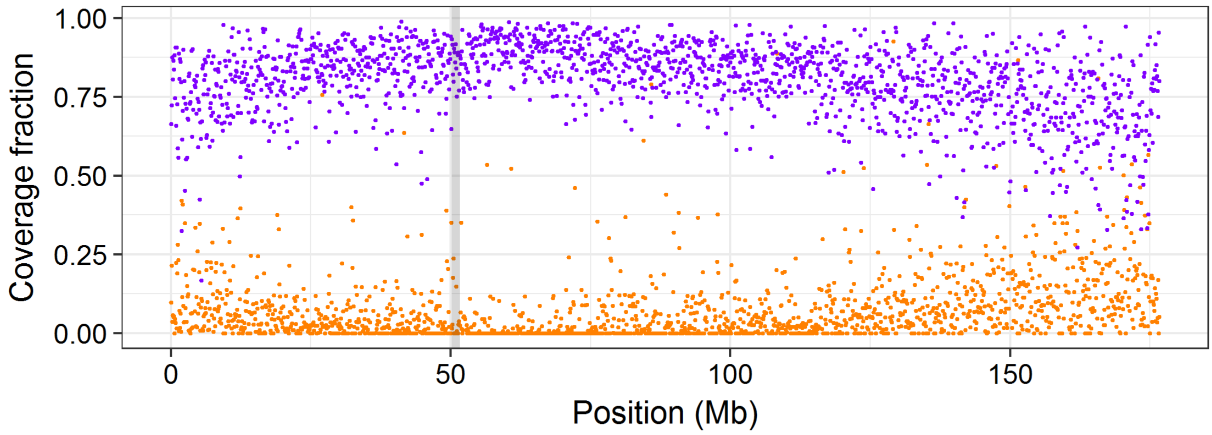


**<u>Supplemental Figure S1 (continued):</u> Methylation lanscape of chromosomes 1 to 10**

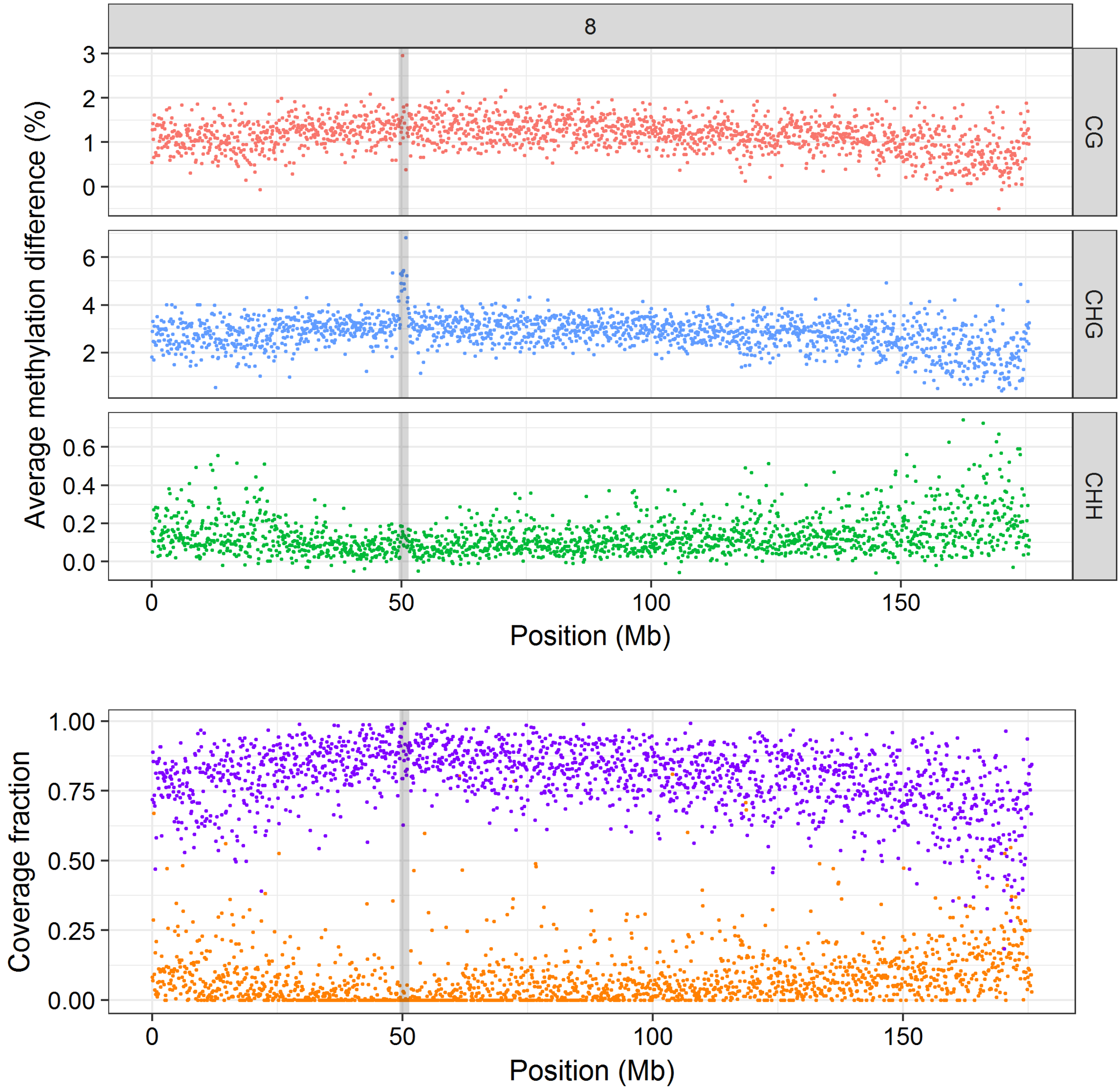


**<u>Supplemental Figure S1 (continued):</u> Methylation lanscape of chromosomes 1 to 10**

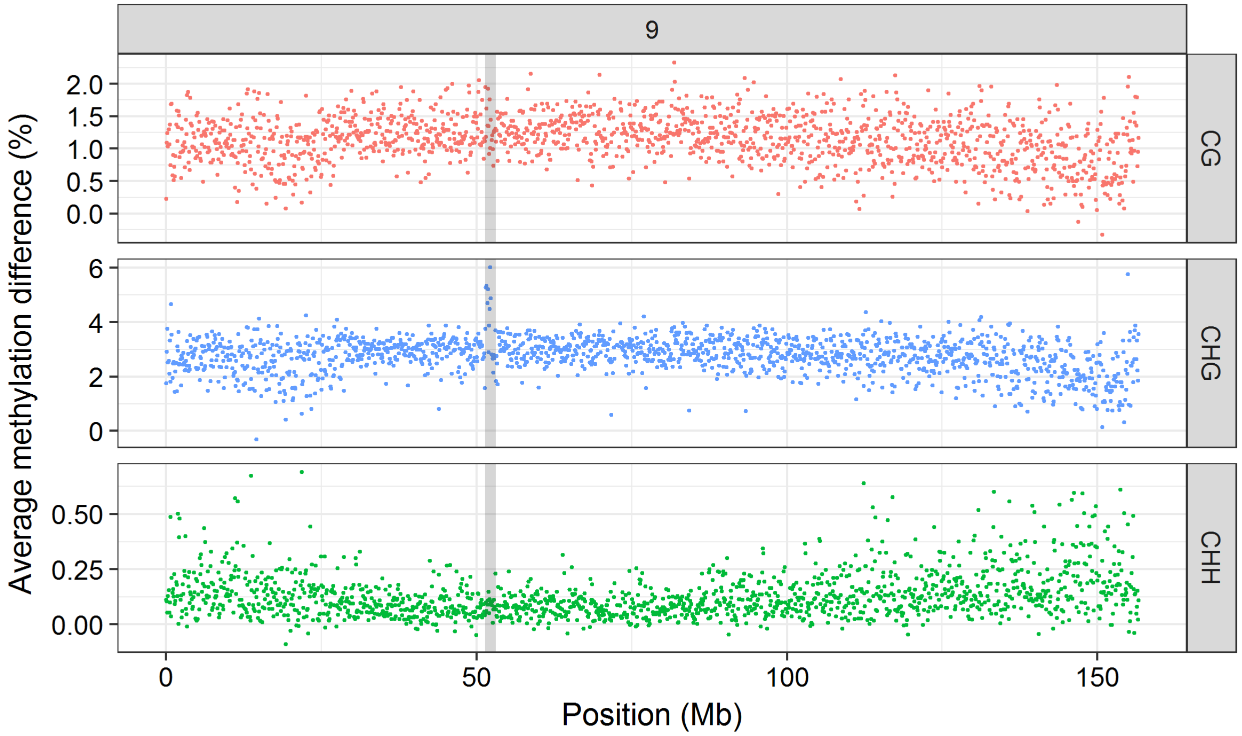


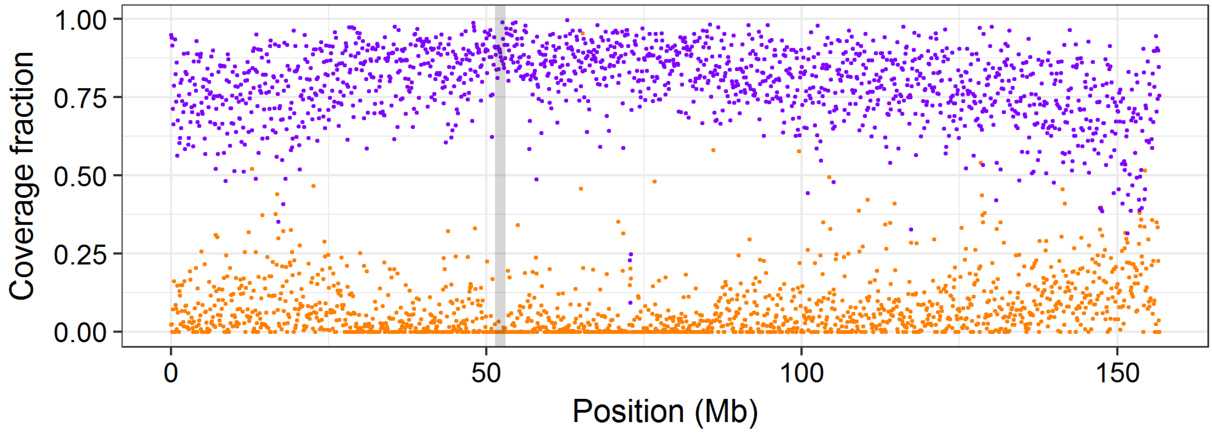


**<u>Supplemental Figure S1 (continued):</u>** **Methylation lanscape of chromosomes 1 to 10**

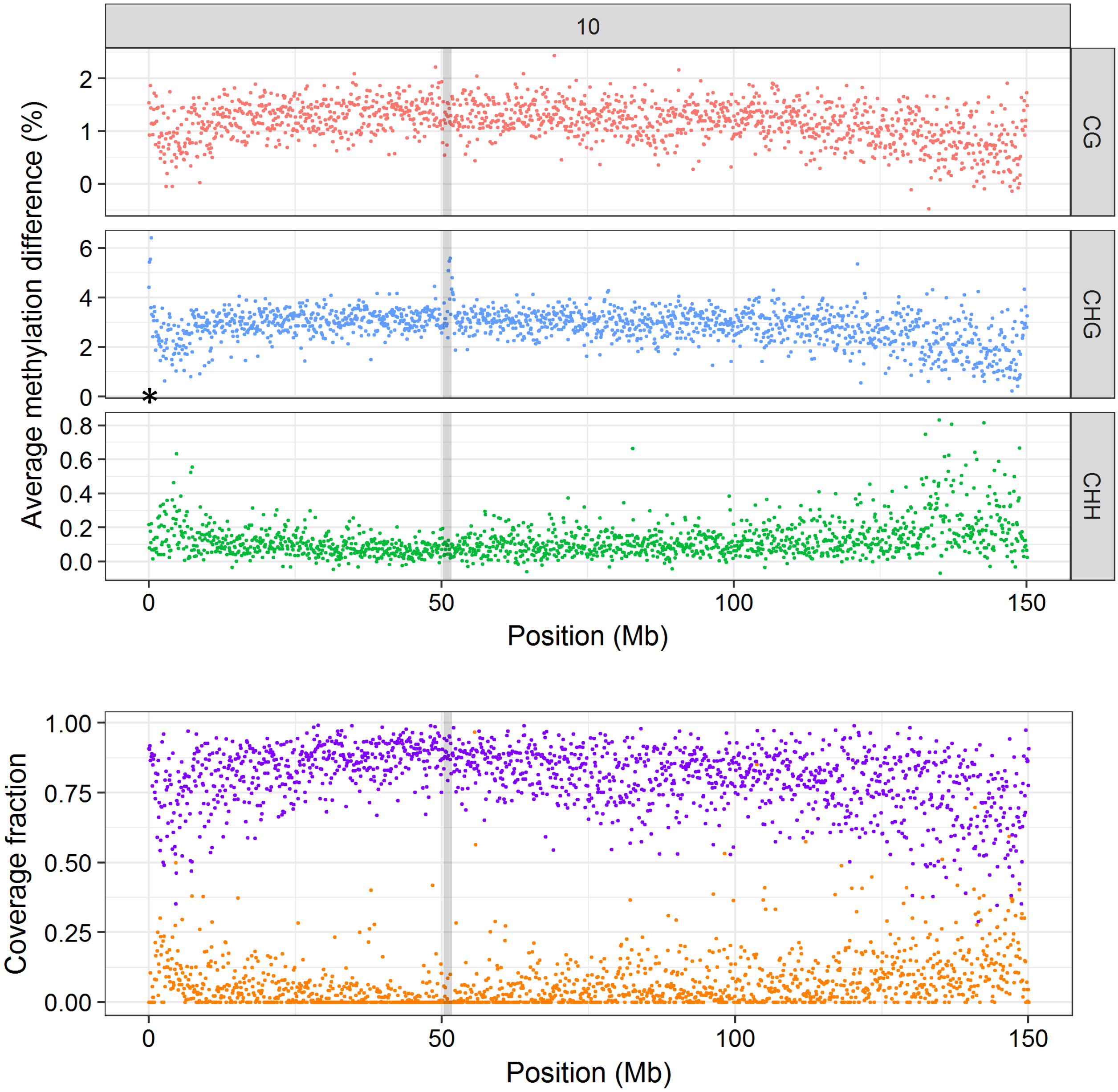


**<u>Supplemental Figure S1 (continued):</u> Methylation lanscape of chromosomes 1 to 10**

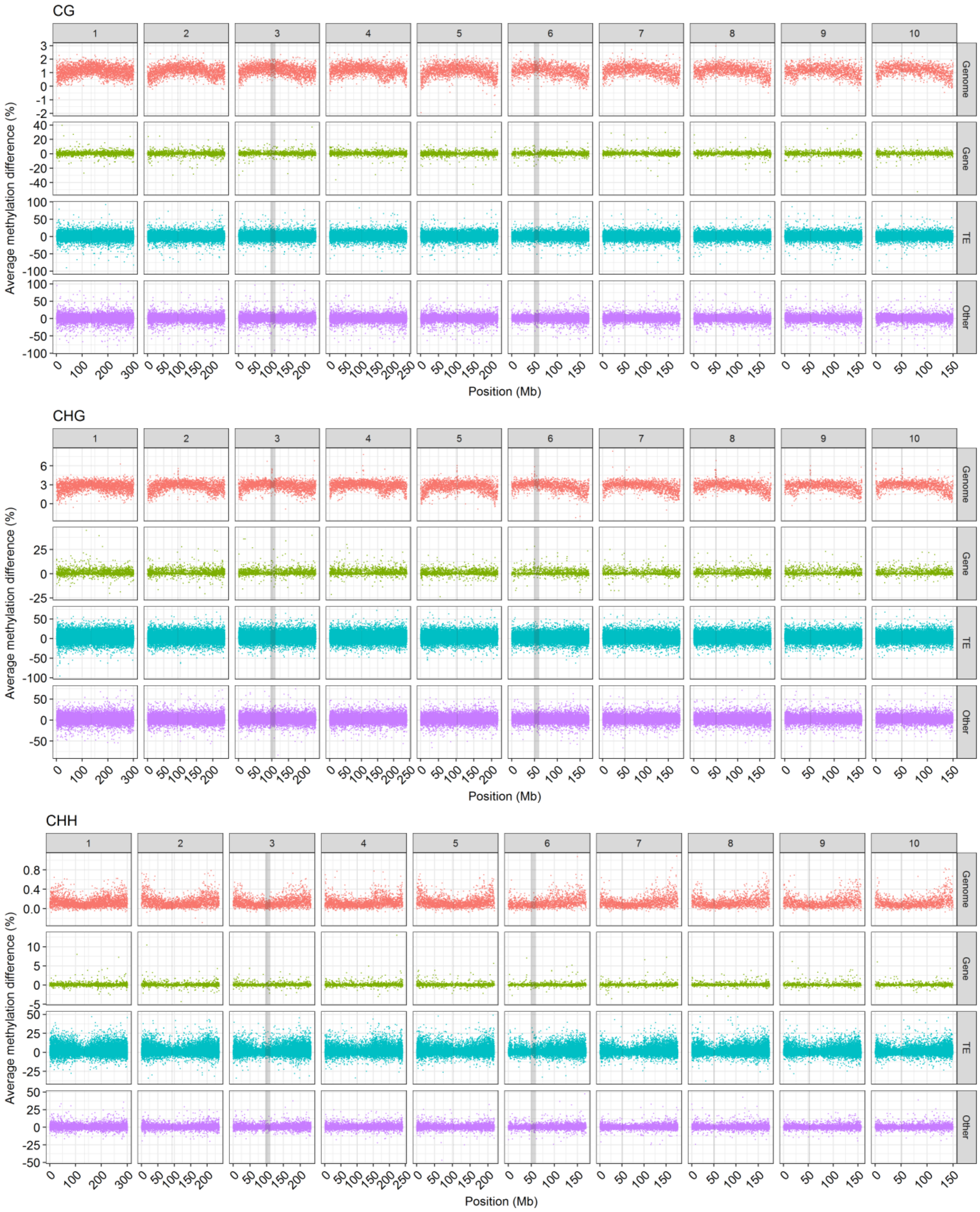


**<u>Supplemental_Figure_S2</u>: Chromosome landscapes of average methylation difference for CG, CHG and CHH cytosine contexts and for the different types of genomic features**

Methylation difference landscape for each category of genomic feature (in line) and each chromosome (in column). Genomic feature type is shown on the right and chromosome numbers are indicated on top. Each dot corresponds to one genomic region. For genome, regions are all of 100 kb. For features, sizes are variable. Centromere positions are highlighted by grey boxes. Cytosine contexts are shown on top of each panel.

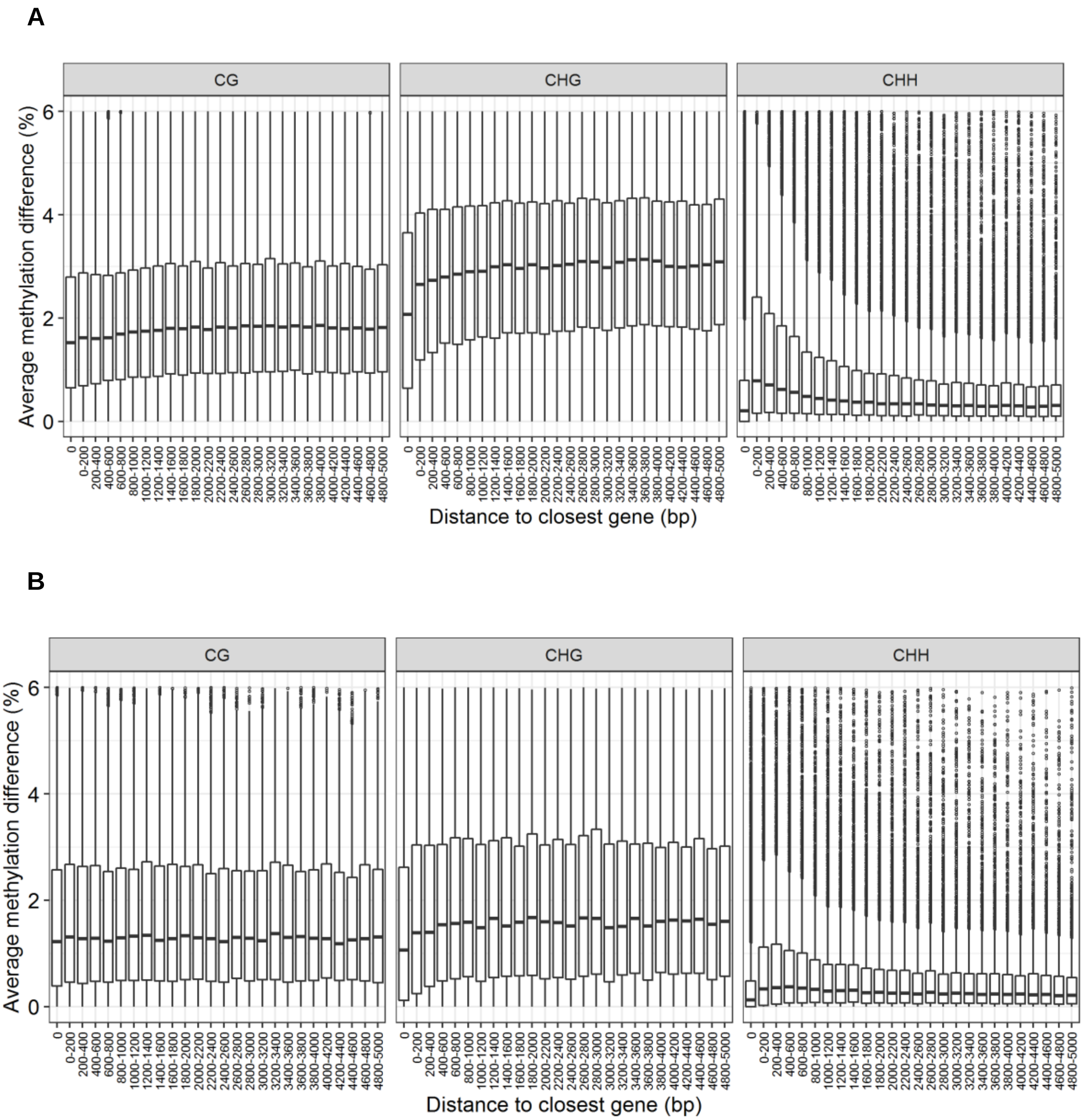


**<u>Supplemental Figure S3</u>: Distribution of methylation difference in TEs for CG, CHG and CHH contexts with respect to distance to closest gene**

Distribution of methylation difference in TEs for CG, CHG and CHH contexts, with respect to distance to closest gene. **A.** Hypermethylated cases. **B.** Hypomethylated cases.

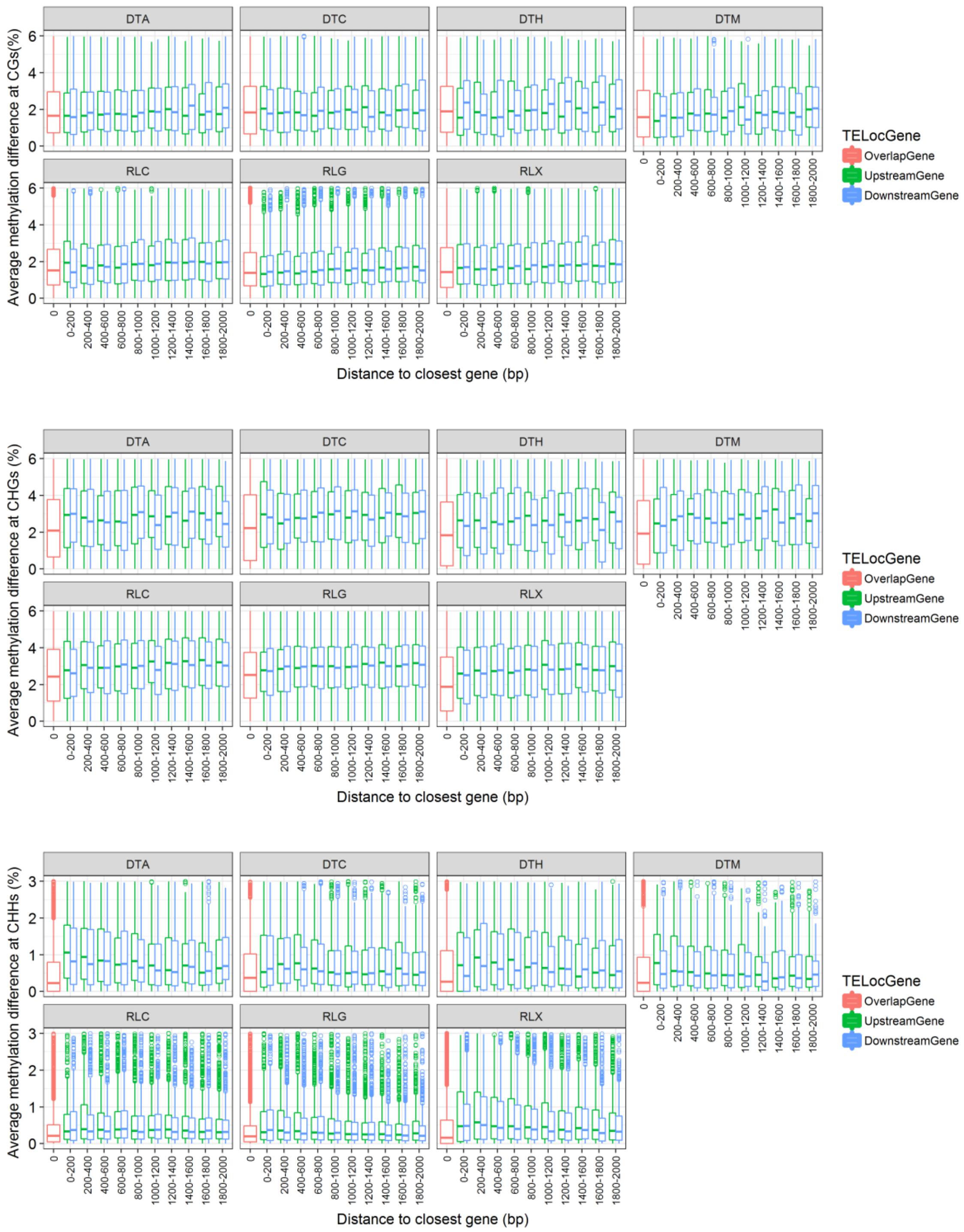


**<u>Supplemental Figure S4:</u> Methylation difference for hypermethylated TEs with respect to gene location and cytosine context**

TEs located upstream (green), downstream (blue), or overlapping (red) genes are shown. Each panel shows one TE class. DTT, RIL and RIX are not represented due to too low sample size.

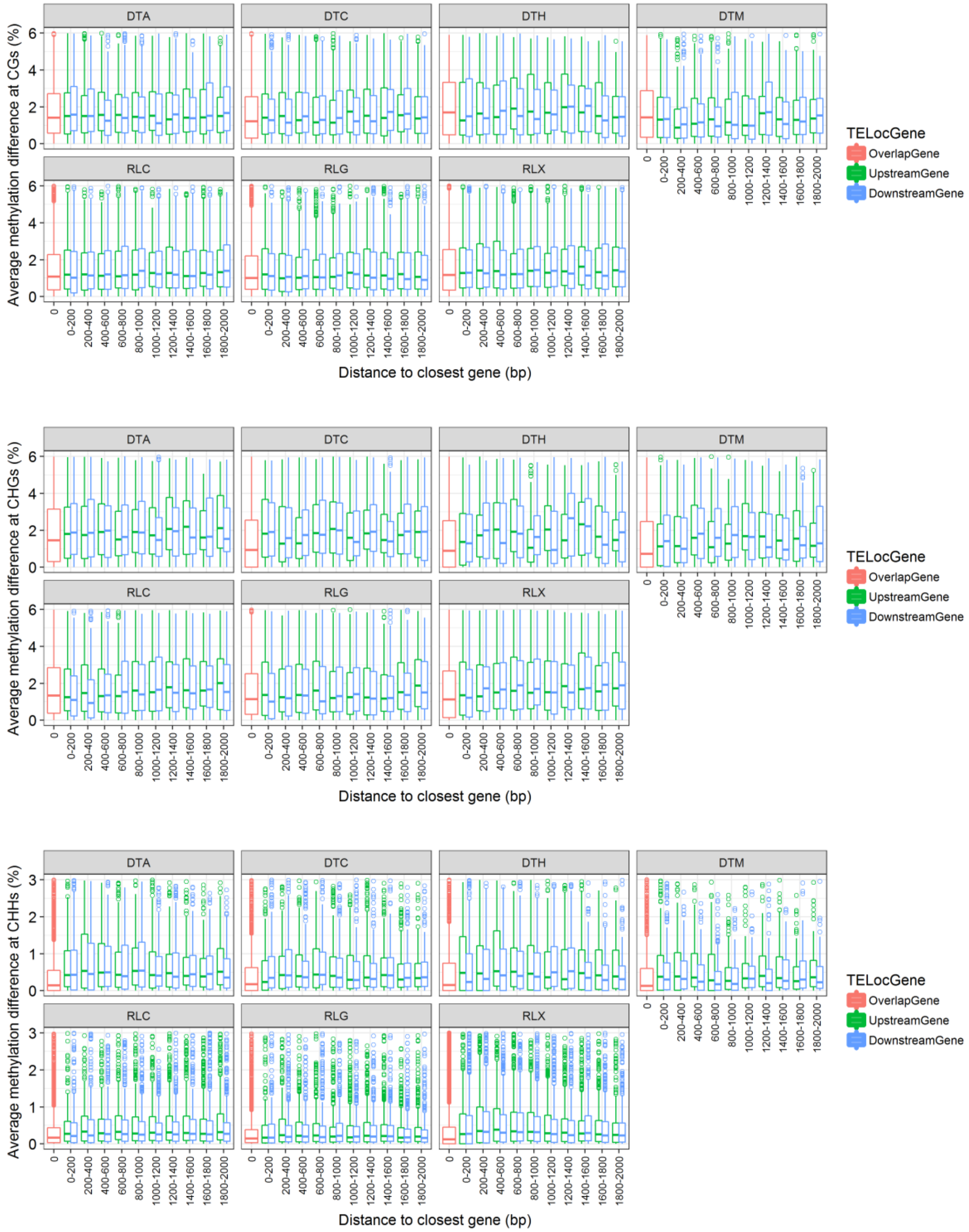


**<u>Supplemental Figure S5:</u> Methylation difference for hypomethylated TEs with respect to gene location and cytosine context**

TEs located upstream (green), downstream (blue), or overlapping (red) genes are shown. Each panel shows one TE class. DTT, RIL and RIX are not represented due to too low sample size.

A

B

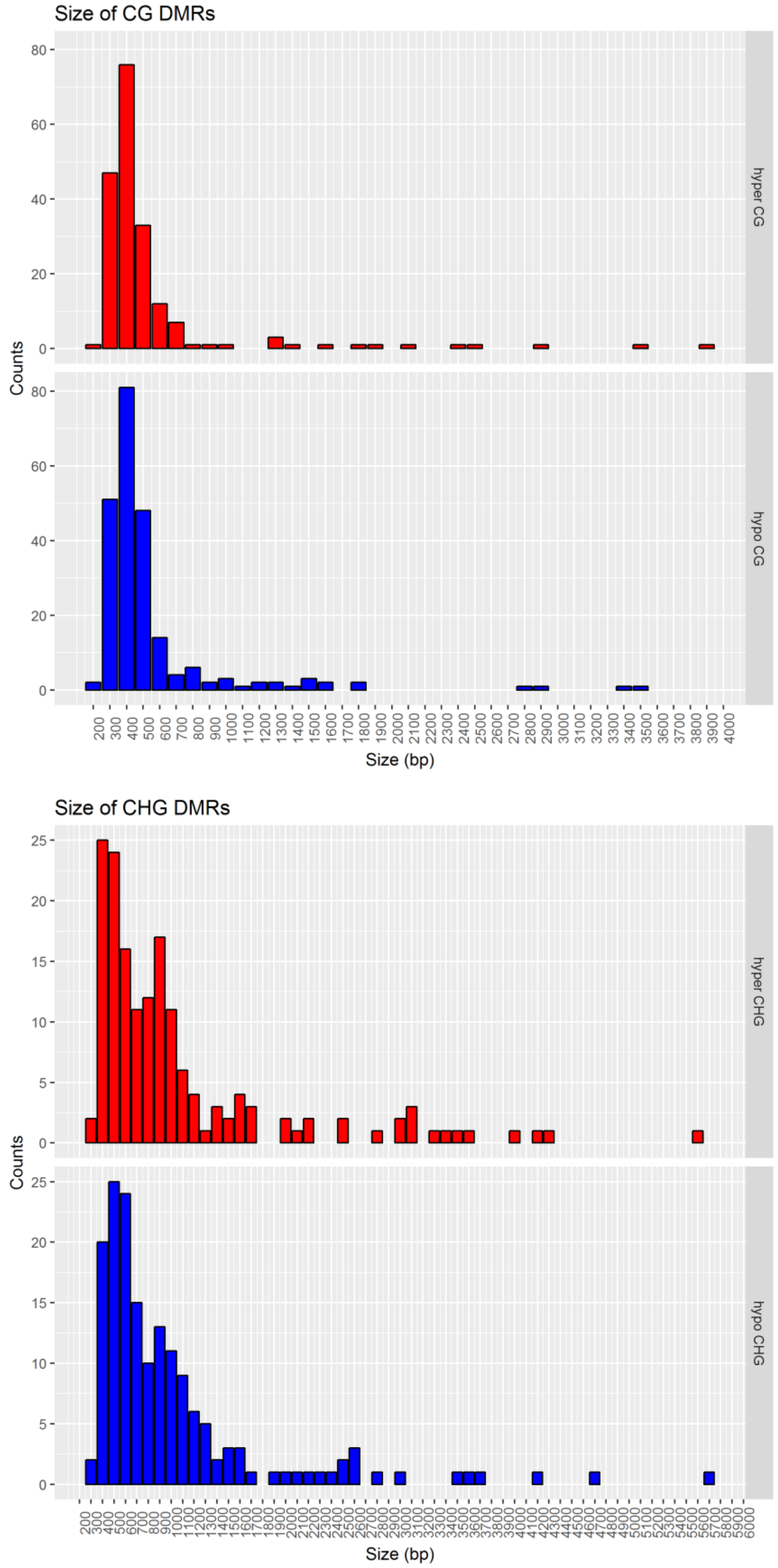


**<u>Supplemental Figure S6:</u> Distribution of DMR size in hyper- (red) and hypomethylated (blue) DMR cases**

**A**. DMRs detected in CG context. **B**. DMRs detected in CHG context.

A

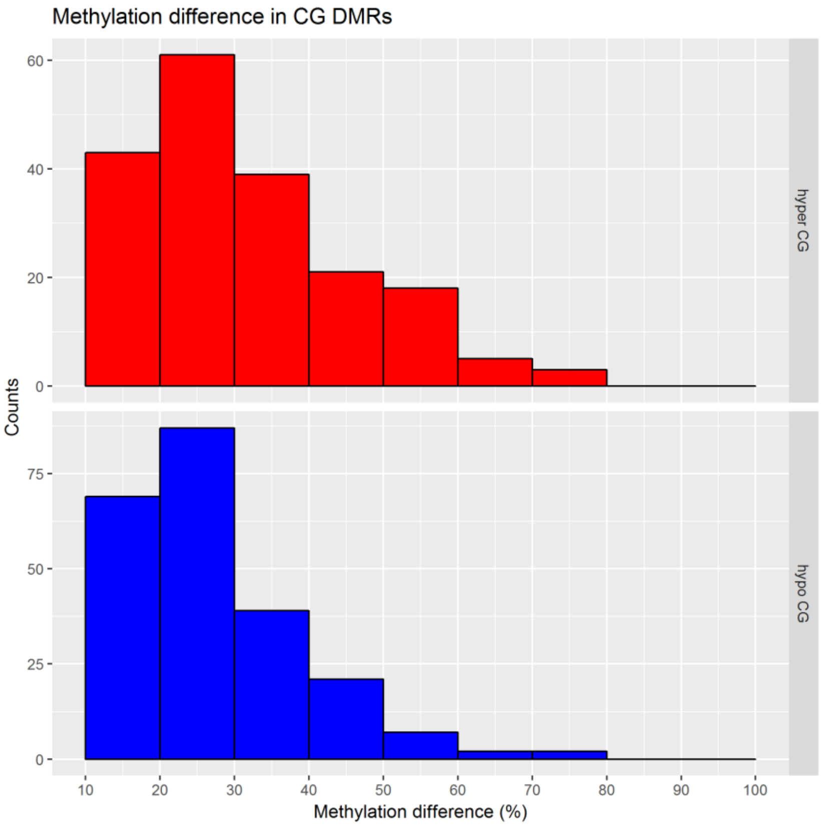


B

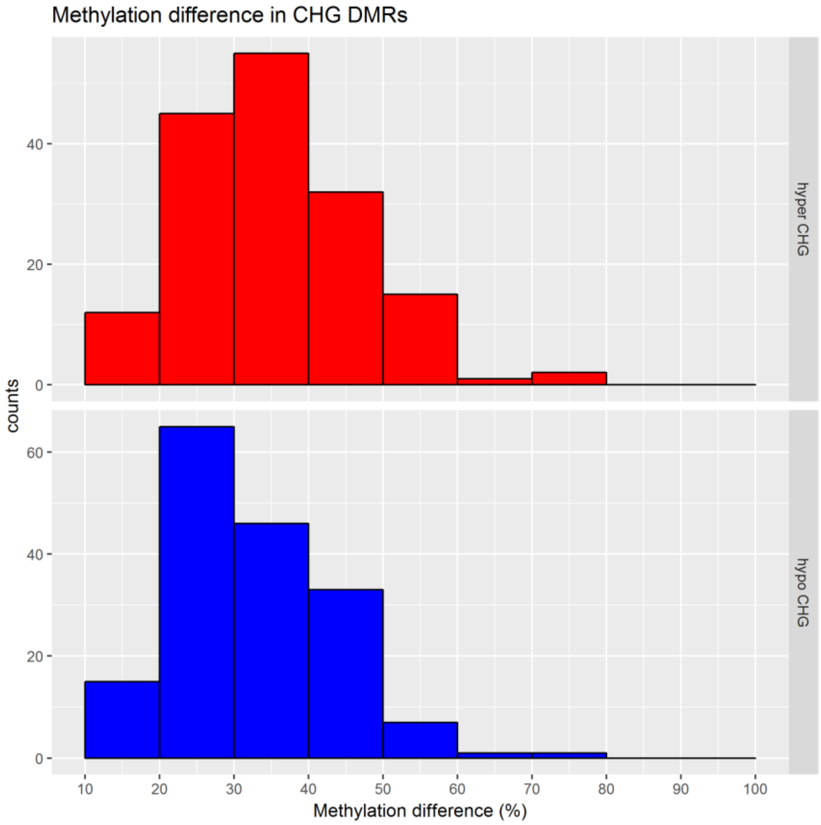


**<u>Supplemental Figure S7:</u> Distribution of average methylation difference in hyper- (red) and hypomethylated (blue) DMR cases**

**A**. DMRs detected in CG context. **B**. DMRs detected in CHG context.

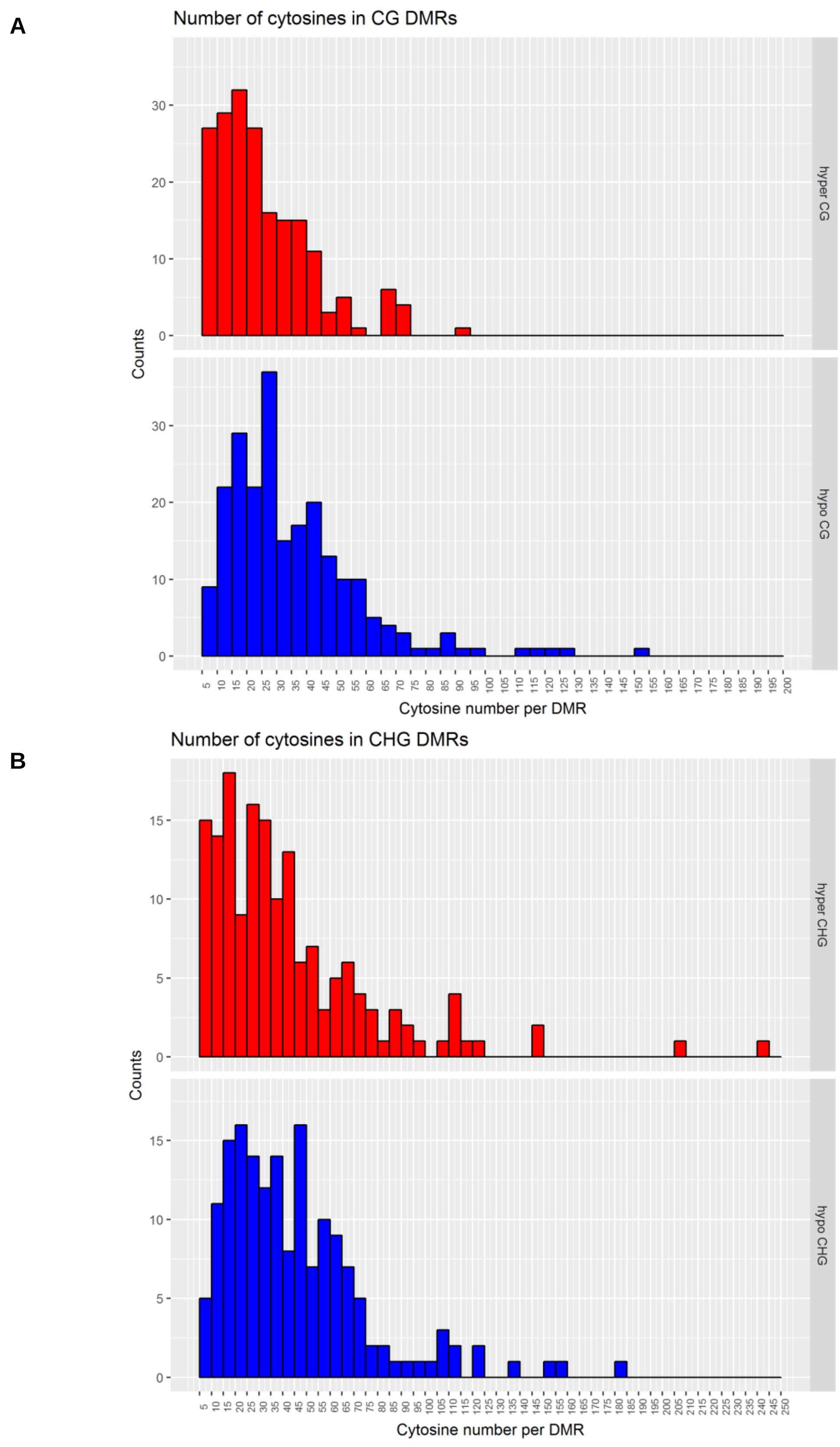


**<u>Supplemental Figure S8:</u> Distribution of cytosine number in hyper- (red) and hypomethylated (blue) DMR cases**

**A**. DMRs detected in CG context. **B**. DMRs detected in CHG context.

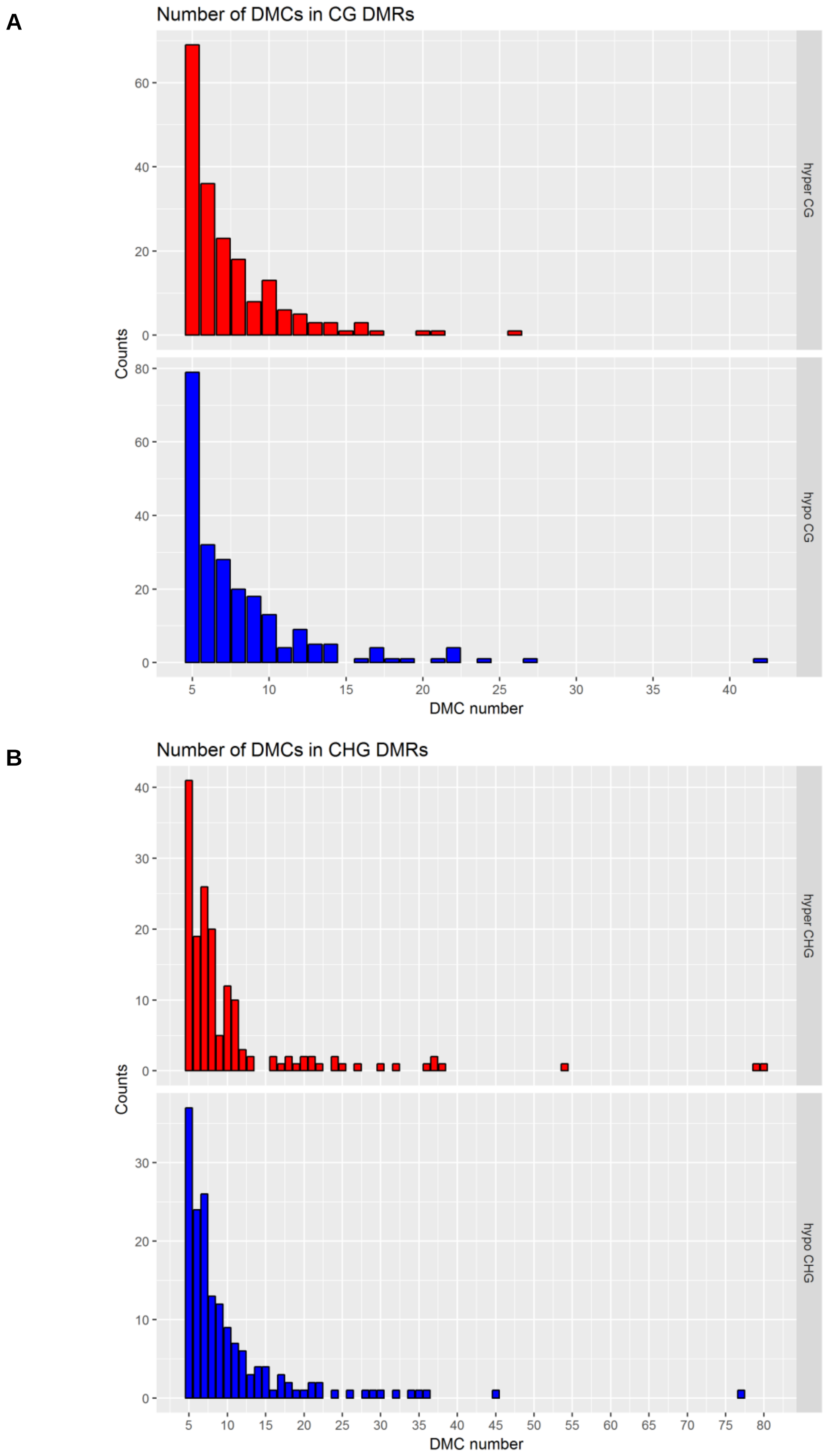


**<u>Supplemental Figure S9:</u> Distribution of DMC number in hyper (red) and hypo (blue) DMR cases**

**A**. DMRs detected in CG context. **B**. DMRs detected in CHG context.

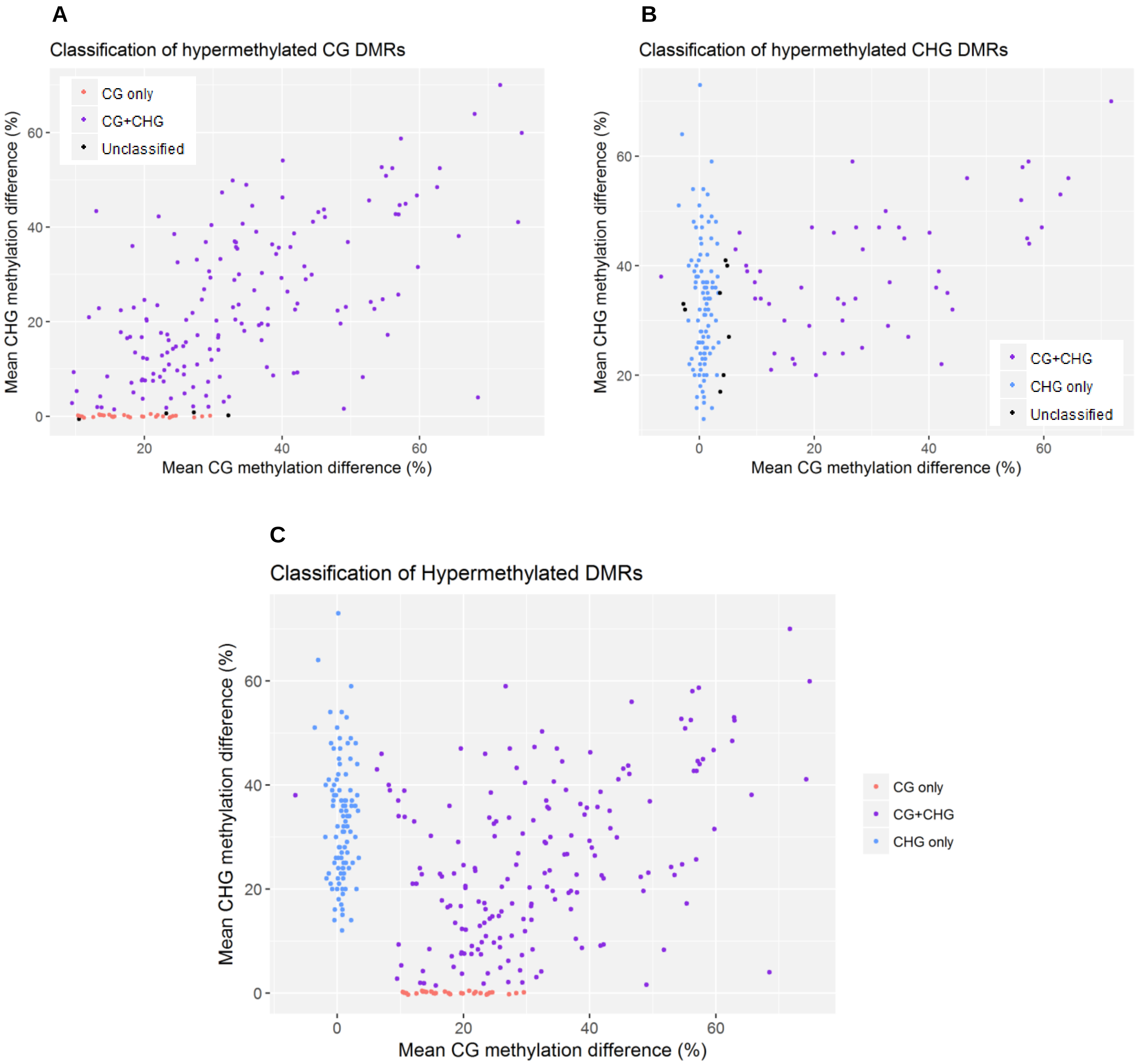


**<u>Supplemental Figure S10:</u> Comparison of average CG and CHG methylation difference between stressed and unstressed plants for hypermethylated DMR cases**

**A**. DMRs detected in CG context. Each dot represents one DMR, and shows corresponding average methylation difference in CG context (x-axis) and in CHG context (y-axis). DMRs are classified as variable in both context (CG+CHG, purple), variable in CG only (red) or unclassified (black). **B**. DMRs detected in CHG context. Each dot represents one DMR, and shows corresponding average methylation difference in CG context (x-axis) and in CHG context (y-axis). DMRs are classified as variable in both context (CG+CHG, purple), variable in CHG only (blue) or unclassified (black). **C**. Final classification of DMRs after removal of unclassified and redundant cases. Each dot represents one DMR, and shows corresponding average methylation difference in CG context (x-axis) and in CHG context (y-axis). DMRs are classified as variable in both context (CG+CHG, purple), variable in CG only (red), or variable in CHG only (blue).

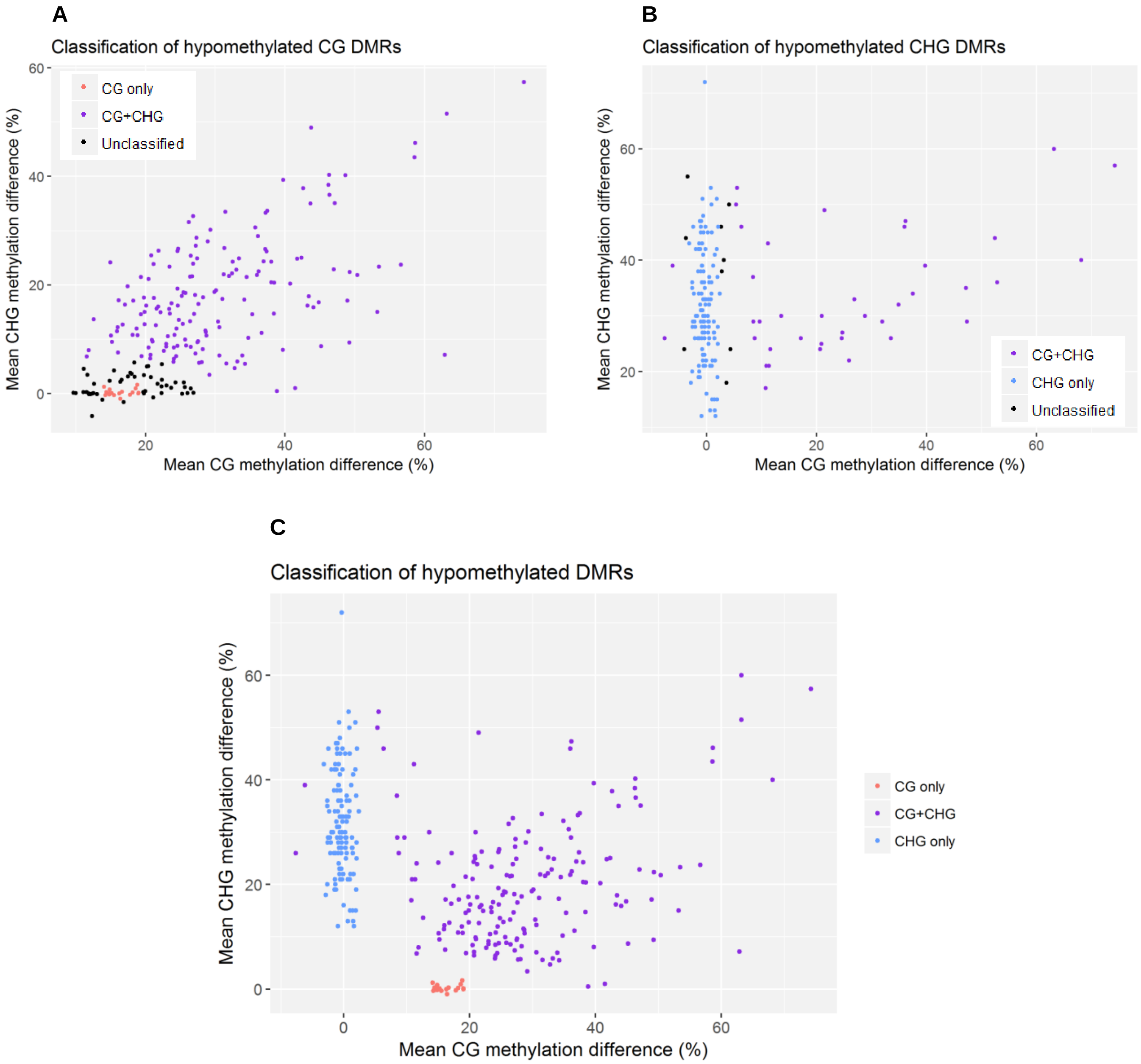


**Supplemental Figure S11: Comparison of average CG and CHG methylation difference between stressed and unstressed plants for hypomethylated DMR cases**

**A**. DMRs detected in CG context. Each dot represents one DMR, and shows corresponding average methylation difference in CG context (x-axis) and in CHG context (y-axis). DMRs are classified as variable in both context (CG+CHG, purple), variable in CG only (red) or unclassified (black). **B**. DMRs detected in CHG context. Each dot represents one DMR, and shows corresponding average methylation difference in CG context (x-axis) and in CHG context (y-axis). DMRs are classified as variable in both context (CG+CHG, purple), variable in CHG only (blue) or unclassified (black). **C**. Final classification of DMRs after removal of unclassified and redundant cases. Each dot represents one DMR, and shows corresponding average methylation difference in CG context (x-axis) and in CHG context (y-axis). DMRs are classified as variable in both context (CG+CHG, purple), variable in CG only (red), or variable in CHG only (blue).

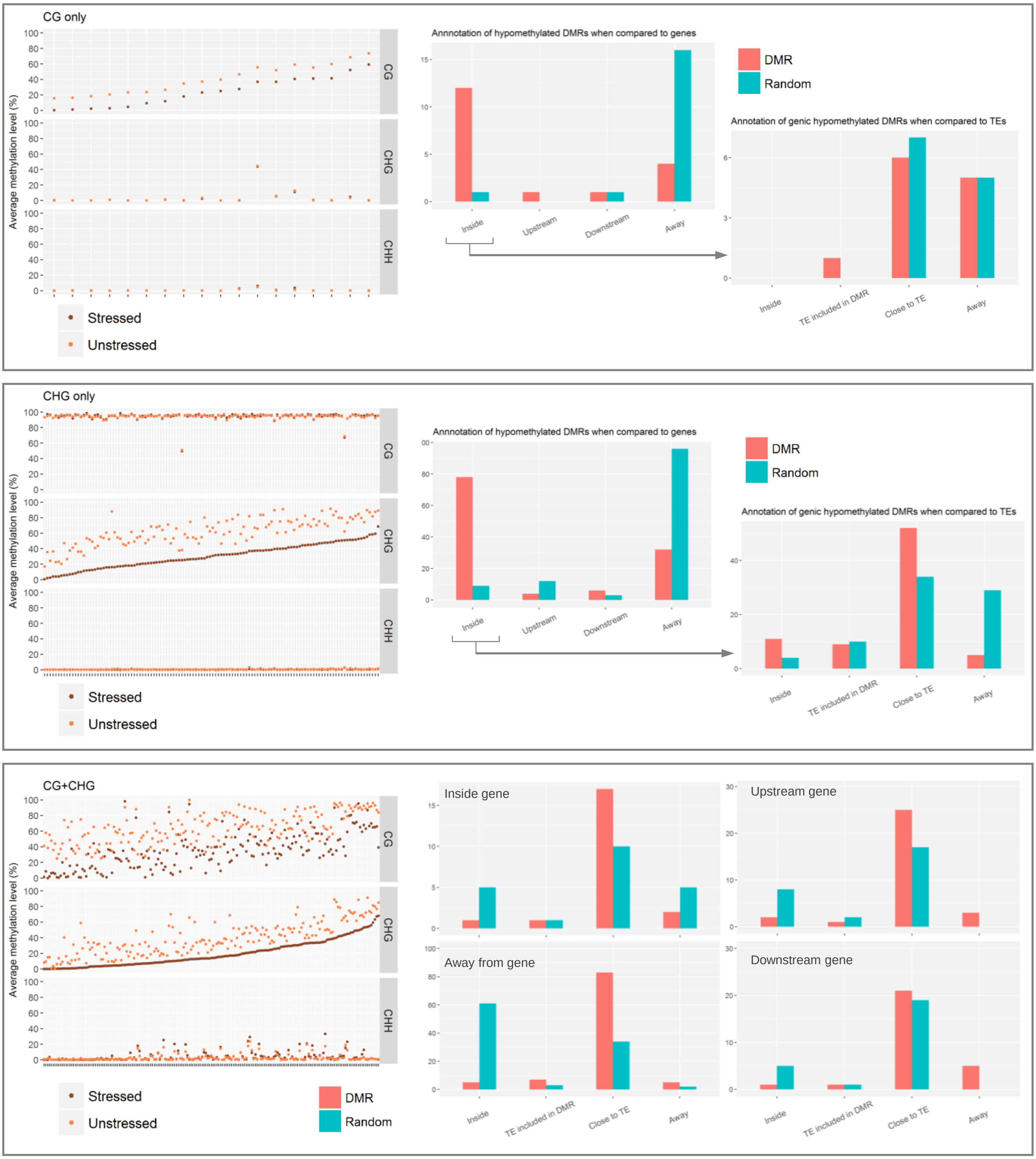


**<u>Supplemental Figure S12:</u> Characteristics of hypomethylated DMRs**

**A.** Caracteristics of CG only DMRs. Left: methylation levels in stressed and unstressed plants. Each position of the x axis corresponds to one DMR, to which two dots correspond, with average values in stressed (brown) and unstressed (orange) plants. Middle: position relative to genes as compared to random regions. For category with largest enrichment (DMRs located inside genes), position relative to TEs is given on the right panel, as compared a random set of regions located inside genes. **B.** Caracteristics of CHG only DMRs. Same legend as A. **C**. Caracteristics of CG+CHG DMRs. Left: same legend as panels A and B. Right four panels: DMR position relative to TEs as compared to random regions, after subdivision of DMRs as compared to distance to genes. Distance thresholds are 2kb and 500bp, for genes and TEs, respectively.

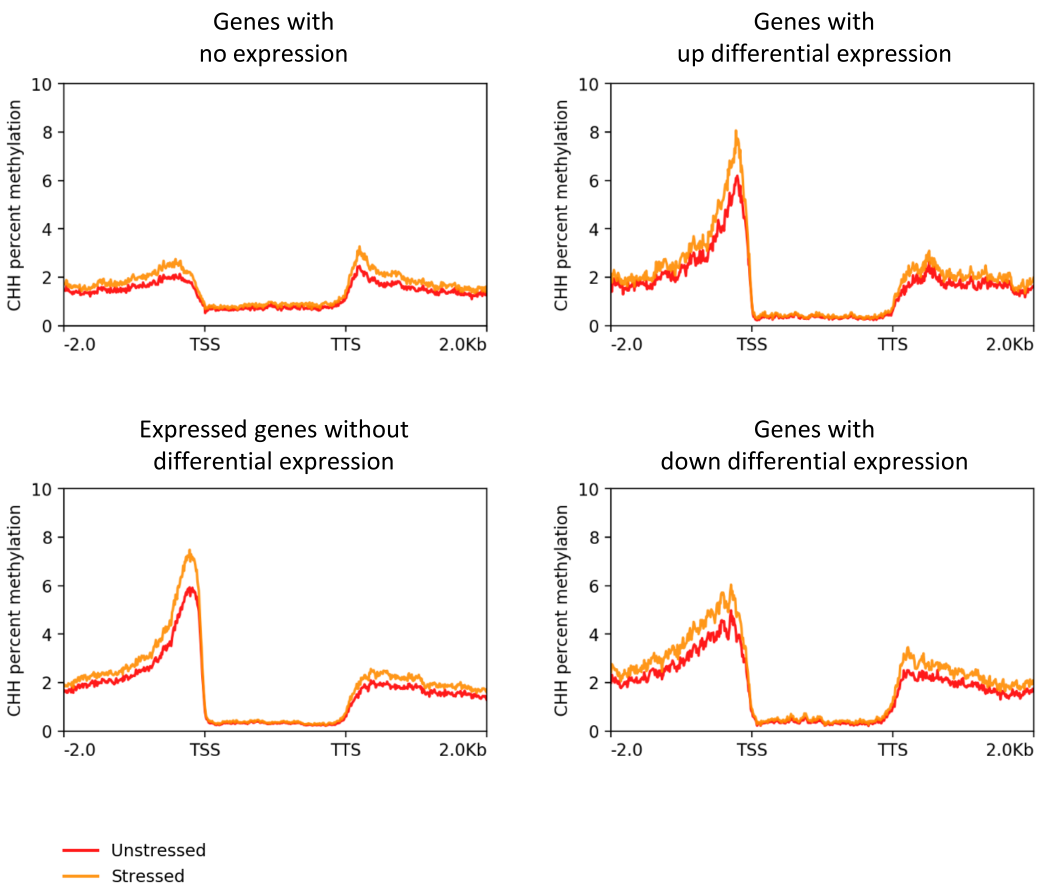


**<u>Supplemental Figure S13</u>: Average CHH methylation levels within genes and in flanking regions**

Comparison of CHH methylation levels in genes and flanking regions for stressed (orange) and unstressed (red) plants. Genes are separated into unexpressed genes, expressed genes without differential expression, genes with up differential expression and genes with down differential expression based on statistical differential analysis. Gene category is shown on top of each panel. Distance to gene Transcription Start Site (TSS) and Transcription Termination Site (TTS) is shown in kb. Gene size is normalized to 2 kb.
